# Interrogating theoretical models of neural computation with emergent property inference

**DOI:** 10.1101/837567

**Authors:** Sean R. Bittner, Agostina Palmigiano, Alex T. Piet, Chunyu A. Duan, Carlos D. Brody, Kenneth D. Miller, John P. Cunningham

## Abstract

A cornerstone of theoretical neuroscience is the circuit model: a system of equations that captures a hypothesized neural mechanism. Such models are valuable when they give rise to an experimentally observed phenomenon – whether behavioral or a pattern of neural activity – and thus can offer insights into neural computation. The operation of these circuits, like all models, critically depends on the choice of model parameters. A key step is then to identify the model parameters consistent with observed phenomena: to solve the inverse problem. In this work, we present a novel technique, emergent property inference (EPI), that brings the modern probabilistic modeling toolkit to theoretical neuroscience. When theorizing circuit models, theoreticians predominantly focus on reproducing computational properties rather than a particular dataset. Our method uses deep neural networks to learn parameter distributions with these computational properties. This methodology is introduced through a motivational example inferring conductance parameters in a circuit model of the stomatogastric ganglion. Then, with recurrent neural networks of increasing size, we show that EPI allows precise control over the behavior of inferred parameters, and that EPI scales better in parameter dimension than alternative techniques. In the remainder of this work, we present novel theoretical findings gained through the examination of complex parametric structure captured by EPI. In a model of primary visual cortex, we discovered how connectivity with multiple inhibitory subtypes shapes variability in the excitatory population. Finally, in a model of superior colliculus, we identified and characterized two distinct regimes of connectivity that facilitate switching between opposite tasks amidst interleaved trials, characterized each regime via insights afforded by EPI, and found conditions where these circuit models reproduce results from optogenetic silencing experiments. Beyond its scientific contribution, this work illustrates the variety of analyses possible once deep learning is harnessed towards solving theoretical inverse problems.

## 2 Introduction

The fundamental practice of theoretical neuroscience is to use a mathematical model to understand neural computation, whether that computation enables perception, action, or some intermediate processing. A neural circuit is systematized with a set of equations – the model – and these equations are motivated by biophysics, neurophysiology, and other conceptual considerations [1–5]. The function of this system is governed by the choice of model *parameters*, which when configured in a particular way, give rise to a measurable signature of a computation. The work of analyzing a model then requires solving the inverse problem: given a computation of interest, how can we reason about the distribution of parameters that give rise to it? The inverse problem is crucial for reasoning about likely parameter values, uniquenesses and degeneracies, and predictions made by the model [6–8].

Ideally, one carefully designs a model and analytically derives how computational properties determine model parameters. Seminal examples of this gold standard include our field’s understanding of memory capacity in associative neural networks [9], chaos and autocorrelation timescales in random neural networks [10], central pattern generation [11], the paradoxical effect [12], and decision making [13]. Unfortunately, as circuit models include more biological realism, theory via analytical derivation becomes intractable. Absent this analysis, statistical inference offers a toolkit by which to solve the inverse problem by identifying, at least approximately, the distribution of parameters that produce computations in a biologically realistic model [14–19].

Statistical inference, of course, requires quantification of the sometimes vague term *computation*. In neuroscience, two perspectives are dominant. First, often we directly use an *exemplar dataset*: a collection of samples that express the computation of interest, this data being gathered either experimentally in the lab or from a computer simulation. Though a natural choice given its connection to experiment [20], some drawbacks exist: these data are well known to have features irrelevant to the computation of interest [21–23], confounding inferences made on such data. Related to this point, use of a conventional dataset encourages conventional data likelihoods or loss functions, which focus on some global metric like squared error or marginal evidence, rather than the computation itself.

Alternatively, researchers often quantify an *emergent property* (EP): a statistic of data that directly quantifies the computation of interest, wherein the dataset is implicit. While such a choice may seem esoteric, it is not: the above “gold standard” examples [9–13] all quantify and focus on some derived feature of the data, rather than the data drawn from the model. An emergent property is of course a dataset by another name, but it suggests different approach to solving the same inverse problem: here we directly specify the desired emergent property – a statistic of data drawn from the model – and the value we wish that property to have, and we set up an optimization program to find the distribution of parameters that produce this computation. This statistical framework is not new: it is intimately connected to the literature on approximate bayesian computation [24–26], parameter sensitivity analyses [27–30], maximum entropy modeling [31–33], and approximate bayesian inference [34, 35]; we detail these connections in Section 5.1.1.

The parameter distributions producing a computation may be curved or multimodal along various parameter axes and combinations. It is by quantifying this complex structure that emergent property inference offers scientific insight. Traditional approximation families (e.g. mean-field or mixture of gaussians) are limited in the distributional structure they may learn. To address such restrictions on expressivity, advances in machine learning have used deep probability distributions as flexible approximating families for such complicated distributions [36, 37] (see Section 5.1.2). However, the adaptation of deep probability distributions to the problem of theoretical circuit analysis requires recent developments in deep learning for constrained optimization [38], and architectural choices for efficient and expressive deep generative modeling [39, 40]. We detail our method, which we call emergent property inference (EPI) in Section 3.2.

Equipped with this method, we demonstrate the capabilities of EPI and present novel theoretical findings from its analysis. First, we show EPI’s ability to handle biologically realistic circuit models using a five-neuron model of the stomatogastric ganglion [41]: a neural circuit whose parametric degeneracy is closely studied [42]. Then, we show EPI’s scalability to high dimensional parameter distributions by inferring connectivities of recurrent neural networks that exhibit stable, yet amplified responses – a hallmark of neural responses throughout the brain [43–45]. In a model of primary visual cortex [46, 47], EPI reveals how the recurrent processing across different neuron-type populations shapes excitatory variability: a finding that we show is analytically intractable. Finally, we investigated the possible connectivities of a superior colliculus model that allow execution of different tasks on interleaved trials [48]. EPI discovered a rich distribution containing two connectivity regimes with different solution classes. We queried the deep probability distribution learned by EPI to produce a mechanistic understanding of neural responses in each regime. Intriguingly, the inferred connectivities of each regime reproduced results from optogenetic inactivation experiments in markedly different ways. These theoretical insights afforded by EPI illustrate the value of deep inference for the interrogation of neural circuit models.

## 3 Results

### 3.1 Motivating emergent property inference of theoretical models

Consideration of the typical workflow of theoretical modeling clarifies the need for emergent property inference. First, one designs or chooses an existing circuit model that, it is hypothesized, captures the computation of interest. To ground this process in a well-known example, consider the stomatogastric ganglion (STG) of crustaceans, a small neural circuit which generates multiple rhythmic muscle activation patterns for digestion [49]. Despite full knowledge of STG connectivity and a precise characterization of its rhythmic pattern generation, biophysical models of the STG have complicated relationships between circuit parameters and computation [15, 42].

A subcircuit model of the STG [41] is shown schematically in Figure 1A. The fast population (f1 and f2) represents the subnetwork generating the pyloric rhythm and the slow population (s1 and s2) represents the subnetwork of the gastric mill rhythm. The two fast neurons mutually inhibit one another, and spike at a greater frequency than the mutually inhibiting slow neurons. The hub neuron couples with either the fast or slow population, or both depending on modulatory conditions. The jagged connections indicate electrical coupling having electrical conductance *g*_el_, smooth connections in the diagram are inhibitory synaptic projections having strength *g*_synA_ onto the hub neuron, and *g*_synB_ = 5nS for mutual inhibitory connections. Note that the behavior of this model will be critically dependent on its parameterization – the choices of conductance parameters **z** = [*g*_el_, *g*_synA_].

**Figure 1:**
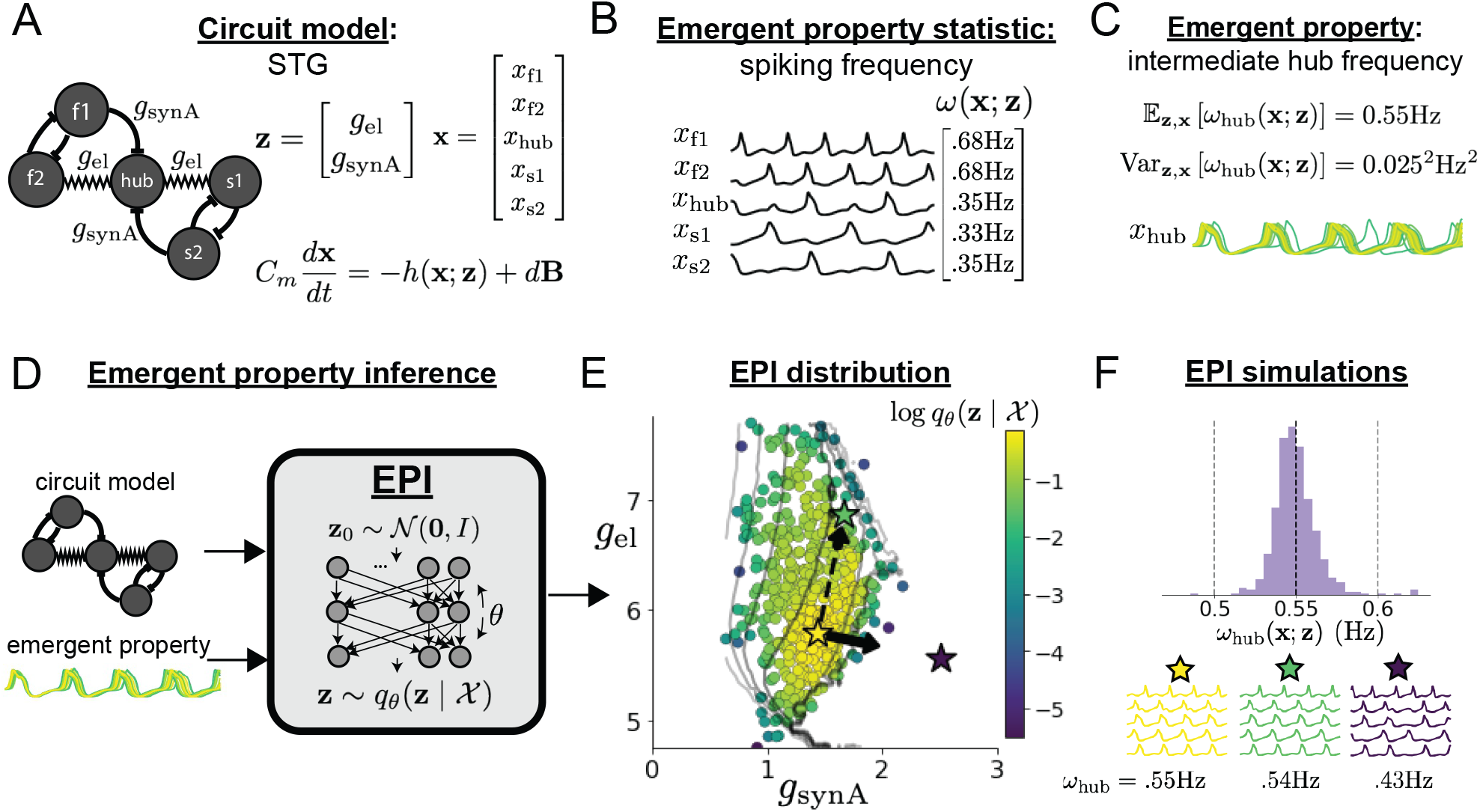
Emergent property inference in the stomatogastric ganglion. **A**. Conductance-based sub-circuit model of the STG. **B**. Spiking frequency *ω*(**x**; **z**) is an emergent property statistic. Simulated at *g*_el_ = 4.5nS and *g*_synA_ = 3nS. **C**. The emergent property of intermediate hub frequency. Simulated activity traces are colored by log probability of generating parameters in the EPI distribution (Panel E). **D**. For a choice of circuit model and emergent property, EPI learns a deep probability distribution of parameters **z**. **E**. The EPI distribution producing intermediate hub frequency. Samples are colored by log probability density. Contours of hub neuron frequency error are shown at levels of .525, .53, … .575 Hz (dark to light gray away from mean). Dimension of sensitivity **v**_1_ (solid arrow) and robustness **v**_2_ (dashed arrow). **F**(Top) The predictions of the EPI distribution. The black and gray dashed lines show the mean and two standard deviations according the emergent property. (Bottom) Simulations at the starred parameter values.

Second, once the model is selected, one must specify what the model should produce. In this STG model, we are concerned with neural spiking frequency, which emerges from the dynamics of the circuit model (Fig. 1B). An emergent property studied by Gutierrez et al. is the hub neuron firing at an intermediate frequency between the intrinsic spiking rates of the fast and slow populations.

This emergent property (EP) is shown in Figure 1C at an average frequency of 0.55Hz. To be precise, we define intermediate hub frequency not strictly as 0.55Hz, but frequencies of moderate deviation from 0.55Hz between the fast (.35Hz) and slow (.68Hz) frequencies.

Third, the model parameters producing the emergent property are inferred. By precisely quantifying the emergent property of interest as a statistical feature of the model, we use emergent property inference (EPI) to condition directly on this emergent property. Before presenting technical details (in the following section), let us understand emergent property inference schematically. EPI (Fig. 1D) takes, as input, the model and the specified emergent property, and as its output, returns the parameter distribution (Fig. 1E). This distribution – represented for clarity as samples from the distribution – is a parameter distribution constrained such that the circuit model produces the emergent property. Once EPI is run, the returned distribution can be used to efficiently generate additional parameter samples. Most importantly, the inferred distribution can be efficiently queried to quantify the parametric structure that it captures. By quantifying the parametric structure governing the emergent property, EPI informs the central question of this inverse problem: what aspects or combinations of model parameters have the desired emergent property?

### 3.2 Emergent property inference via deep generative models

EPI formalizes the three-step procedure of the previous section with deep probability distributions [36, 37]. First, as is typical, we consider the model as a coupled set of noisy differential equations. In this STG example, the model activity (or state) **x** = [*x*_f1_, *x*_f2_, *x*_hub_, *x*_s1_, *x*_s2_] is the membrane potential for each neuron, which evolves according to the biophysical conductance-based equation:

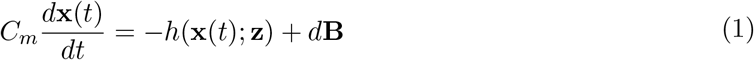

where *C_m_*=1nF, and **h** is a sum of the leak, calcium, potassium, hyperpolarization, electrical, and synaptic currents, all of which have their own complicated dependence on activity **x** and parameters **z** = [*g*_el_, *g*_synA_], and *d***B** is white gaussian noise [41] (see Section 5.2.1 for more detail).

Second, we determine that our model should produce the emergent property of “intermediate hub frequency” (Figure 1C). We stipulate that the hub neuron’s spiking frequency – denoted by statistic *ω*_hub_(**x**) – is close to a frequency of 0.55Hz, between that of the slow and fast frequencies. Mathematically, we define this emergent property with two constraints: that the mean hub frequency is 0.55Hz,

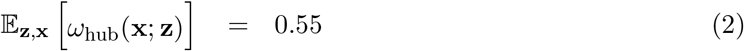

and that the variance of the hub frequency is moderate

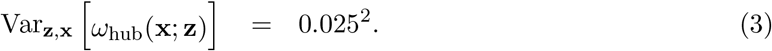

In the emergent property of intermediate hub frequency, the statistic of hub neuron frequency is an expectation over the distribution of parameters **z** and the distribution of the data **x** that those parameters produce. We define the emergent property 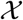 as the collection of these two constraints. In general, an emergent property is a collection of constraints on statistical moments that together define the computation of interest.

Third, we perform emergent property inference: we find a distribution over parameter configurations **z** of models that produce the emergent property; in other words, they satisfy the constraints introduced in Equations 2 and 3. This distribution will be chosen from a family of probability distributions 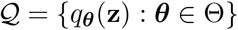, defined by a deep neural network [36, 37] (Figure 1D, EPI box). Deep probability distributions map a simple random variable **z**_0_ (e.g. an isotropic gaussian) through a deep neural network with weights and biases ***θ*** to parameters **z** = *g_**θ**_*(**z**_0_) of a suitably complicated distribution (see Section 5.1.2 for more details). Many distributions in 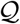 will respect the emergent property constraints, so we select the most random (highest entropy) distribution, which also means this approach is equivalent to bayesian variational inference (see Section 5.1.6). In EPI optimization, stochastic gradient steps in ***θ*** are taken such that entropy is maximized, and the emergent property 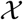 is produced (see Section 5.1). We then denote the inferred EPI distribution as 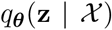, since the structure of the learned parameter distribution is determined by weights and biases ***θ***, and this distribution is conditioned upon emergent property 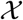.

The structure of the inferred parameter distributions of EPI can be analyzed to reveal key information about how the circuit model produces the emergent property. As probability in the EPI distribution decreases away from the mode of 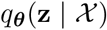 (Fig. 1E yellow star), the emergent property deteriorates. Perturbing **z** along a dimension in which 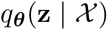 changes little will not disturb the emergent property, making this parameter combination *robust* with respect to the emergent property. In contrast, if **z** is perturbed along a dimension with strongly decreasing 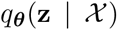, that parameter combination is deemed *sensitive* [27, 30]. By querying the second order derivative (Hessian) of 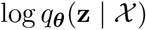 at a mode, we can quantitatively identify how sensitive (or robust) each eigenvector is by its eigenvalue; the more negative, the more sensitive and the closer to zero, the more robust (see Section 5.2.4). Indeed, samples equidistant from the mode along these dimensions of sensitivity (**v**_1_, smaller eigenvalue) and robustness (**v**_2_, greater eigenvalue) (Fig. 1E, arrows) agree with error contours (Fig. 1E contours) and have diminished or preserved hub frequency, respectively (Fig. 1F activity traces). The directionality of **v**_2_ suggests that changes in conductance along this parameter combination will most preserve hub neuron firing between the intrinsic rates of the pyloric and gastric mill rhythms. Importantly and unlike alternative techniques, once an EPI distribution has been learned, the modes and Hessians of the distribution can be measured with trivial computation (see Section 5.1.2).

In the following sections, we demonstrate EPI on three neural circuit models across ranges of biological realism, neural system function, and network scale. First, we demonstrate the superior scalability of EPI compared to alternative techniques by inferring high-dimensional distributions of recurrent neural network connectivities that exhibit amplified, yet stable responses. Next, in a model of primary visual cortex [46,47], we show how EPI discovers parametric degeneracy, revealing how input variability across neuron types affects the excitatory population. Finally, in a model of superior colliculus [48], we used EPI to capture multiple parametric regimes of task switching, and queried the dimensions of parameter sensitivity to characterize each regime.

### 3.3 Scaling inference of recurrent neural network connectivity with EPI

To understand how EPI scales in comparison to existing techniques, we consider recurrent neural networks (RNNs). Transient amplification is a hallmark of neural activity throughout cortex, and is often thought to be intrinsically generated by recurrent connectivity in the responding cortical area [43–45]. It has been shown that to generate such amplified, yet stabilized responses, the connectivity of RNNs must be non-normal [43, 50], and satisfy additional constraints [51]. In theoretical neuroscience, RNNs are optimized and then examined to show how dynamical systems could execute a given computation [52, 53], but such biologically realistic constraints on connectivity [43, 50, 51] are ignored for simplicity or because constrained optimization is difficult. In general, access to distributions of connectivity that produce theoretical criteria like stable amplification, chaotic fluctuations [10], or low tangling [54] would add scientific value to existing research with RNNs. Here, we use EPI to learn RNN connectivities producing stable amplification, and demonstrate the superior scalability and efficiency of EPI to alternative approaches.

We consider a rank-2 RNN with *N* neurons having connectivity *W* = *UV*^⊤^ and dynamics

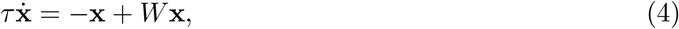

where 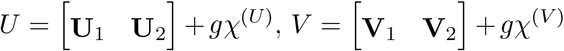, **U**_1_ **U**_2_, **V**_1_, **V**_2_ ∈ [−1, 1]^*N*^, and 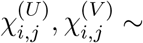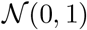. We infer connectivity parameters **z** = [**U**_1_, **U**_2_, **V**_1_, **V**_2_] that produce stable amplification.

Two conditions are necessary and sufficient for RNNs to exhibit stable amplification [51]: real(*λ*_1_) < 1 and 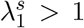, where *λ*_1_ is the eigenvalue of *W* with greatest real part and *λ^s^* is the maximum eigenvalue of 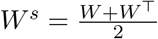. RNNs with real(*λ*_1_) = 0.5 ± 0.5 and 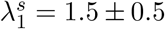 will be stable with modest decay rate (real(*λ*_1_) close to its upper bound of 1) and exhibit modest amplification (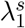 close to its lower bound of 1). EPI can naturally condition on this emergent property

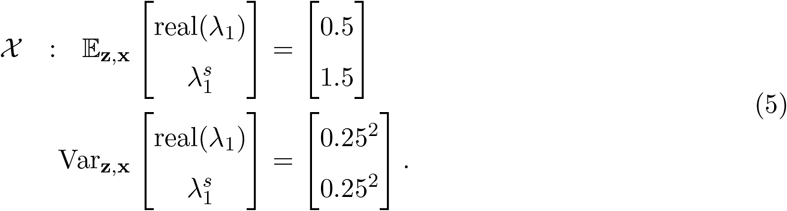

Variance constraints predicate that the majority of the distribution (within two standard deviations) are within the specified ranges.

For comparison, we infer the parameters **z** likely to produce stable amplification using two alternative simulation-based inference approaches. Sequential Monte Carlo approximate bayesian computation (SMC-ABC) [26] is a rejection sampling approach that uses SMC techniques to improve efficiency, and sequential neural posterior estimation (SNPE) [35] approximates posteriors with deep probability distributions (see Section 5.1.1). Unlike EPI, these statistical inference techniques do not constrain the predictions of the inferred distribution, so they were run by conditioning on an exemplar dataset **x**_0_ = ***μ***, following standard practice with these methods [26, 35]. To compare the efficiency of these different techniques, we measured the time and number of simulations necessary for the distance of the predictive mean to be less than 0.5 from ***μ*** = **x**_0_ (see Section 5.3).

As the number of neurons *N* in the RNN, and thus the dimension of the parameter space **z** ∈ [−1, 1]^4*N*^, is scaled, we see that EPI converges at greater speed and at greater dimension than SMC-ABC and SNPE (Fig. 2A). It also becomes most efficient to use EPI in terms of simulation count at *N* = 50 (Fig. 2B). It is well known that ABC techniques struggle in parameter spaces of modest dimension [55], yet we were careful to assess the scalability of SNPE, which is a more closely related methodology to EPI. Between EPI and SNPE, we closely controlled the number of parameters in deep probability distributions by dimensionality (Fig. S5), and tested more aggressive SNPE hyperparameter choices when SNPE failed to converge (Fig. S6). In this analysis, we see that deep inference techniques EPI and SNPE are far more amenable to inference of high dimensional RNN connectivities than rejection sampling techniques like SMC-ABC, and that EPI outperforms SNPE in both wall time (elapsed real time) and simulation count.

**Figure 2:**
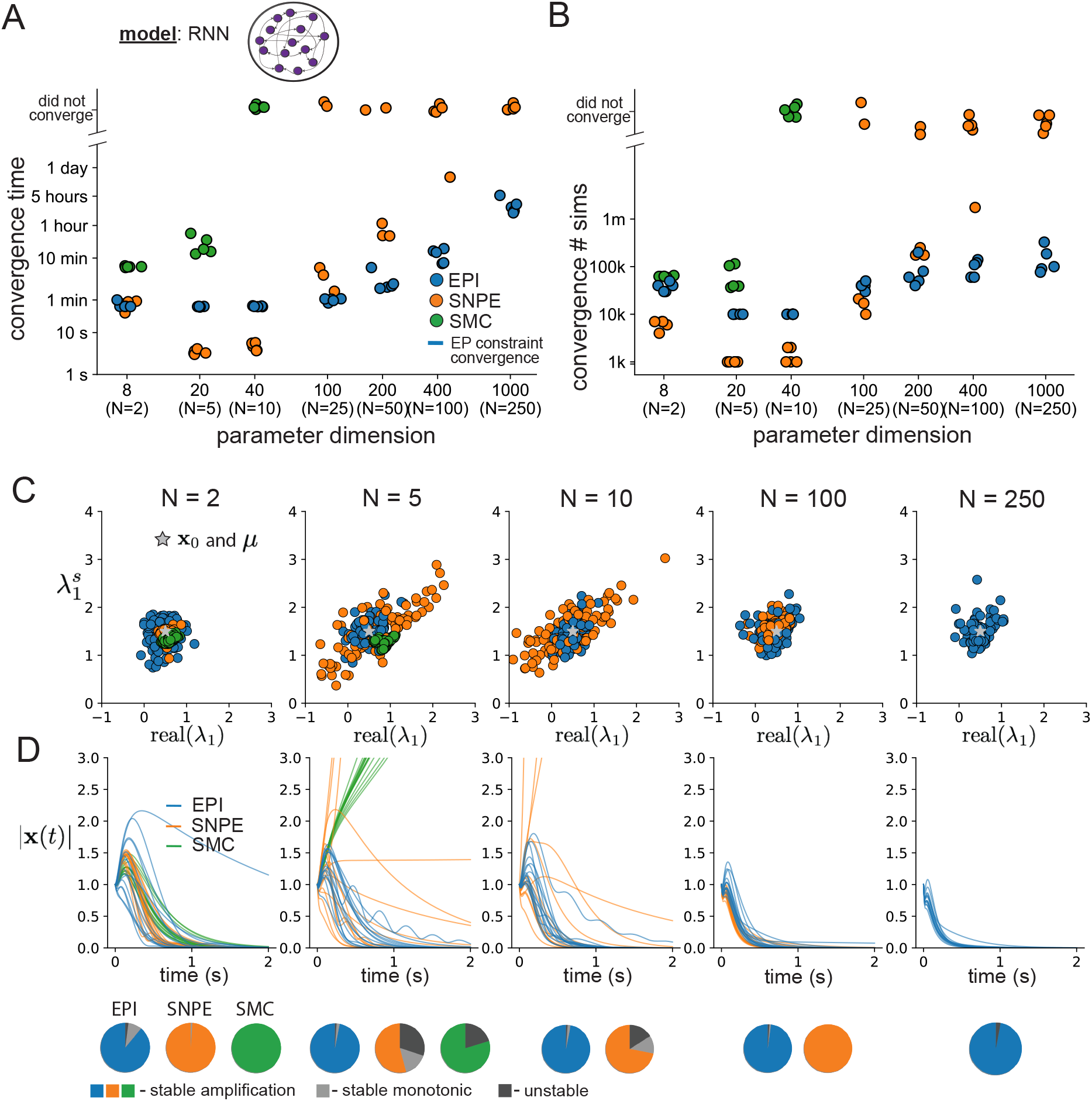
**A**. Wall time of EPI (blue), SNPE (orange), and SMC-ABC (green) to converge on RNN connectivities producing stable amplification. Each dot shows convergence time for an individual random seed. For reference, the mean wall time for EPI to achieve its full constraint convergence (means and variances) is shown (blue line). **B**. Simulation count of each algorithm to achieve convergence. Same conventions as A. **C**. The predictive distributions of connectivities inferred by EPI (blue), SNPE (orange), and SMC-ABC (green), with reference to **x**_0_ = ***μ*** (gray star). **D**. Simulations of networks inferred by each method (*τ* = 100*ms*). Each trace (15 per algorithm) corresponds to simulation of one *z*. (Below) Ratio of obtained samples producing stable amplification, stable monotonic decay, and instability.

No matter the number of neurons, EPI always produces connectivity distributions with mean and variance of real(*λ*_1_) and 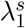 according to 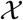 (Fig. 2C, blue). For the dimensionalities in which SMC-ABC is tractable, the inferred parameters are concentrated and offset from the exemplar dataset **x**_0_ (Fig. 2C, green). When using SNPE, the predictions of the inferred parameters are highly concentrated at some RNN sizes and widely varied in others (Fig. 2C, orange). We see these properties reflected in simulations from the inferred distributions: EPI produces a consistent variety of stable, amplified activity norms |**x**(*t*)|, SMC-ABC produces a limited variety of responses, and the changing variety of responses from SNPE emphasizes the control of EPI on parameter predictions (Fig. 2D). Even for moderate neuron counts, the predictions of the inferred distribution of SNPE are highly dependent on *N* and *g*, while EPI maintains the emergent property across choices of RNN (see Section 5.3.5).

To understand these differences, note that EPI outperforms SNPE in high dimensions by using gradient information (from 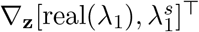). This choice agrees with recent speculation that such gradient information could improve the efficiency of simulation-based inference techniques [56], as well as reflecting the classic tradeoff between gradient-based and sampling-based estimators (scaling and speed versus generality). Since gradients of the emergent property are necessary in EPI optimization, gradient tractability is a key criteria when determining the suitability of a simulation-based inference technique. If the emergent property gradient is efficiently calculated, EPI is a clear choice for inferring high dimensional parameter distributions. In the next two sections, we use EPI for novel scientific insight by examining the structure of inferred distributions.

### 3.4 EPI reveals how recurrence with multiple inhibitory subtypes governs excitatory variability in a V1 model

Dynamical models of excitatory (E) and inhibitory (I) populations with supralinear input-output function have succeeded in explaining a host of experimentally documented phenomena in primary visual cortex (V1). In a regime characterized by inhibitory stabilization of strong recurrent excitation, these models give rise to paradoxical responses [12], selective amplification [43, 50], surround suppression [57] and normalization [58]. Recent theoretical work [59] shows that stabilized E-I models reproduce the effect of variability suppression [60]. Furthermore, experimental evidence shows that inhibition is composed of distinct elements – parvalbumin (P), somatostatin (S), VIP (V) – composing 80% of GABAergic interneurons in V1 [61–63], and that these inhibitory cell types follow specific connectivity patterns (Fig. 3A) [64]. Here, we use EPI on a model of V1 with biologically realistic connectivity to show how the structure of input across neuron types affects the variability of the excitatory population – the population largely responsible for projecting to other brain areas [65].

**Figure 3:**
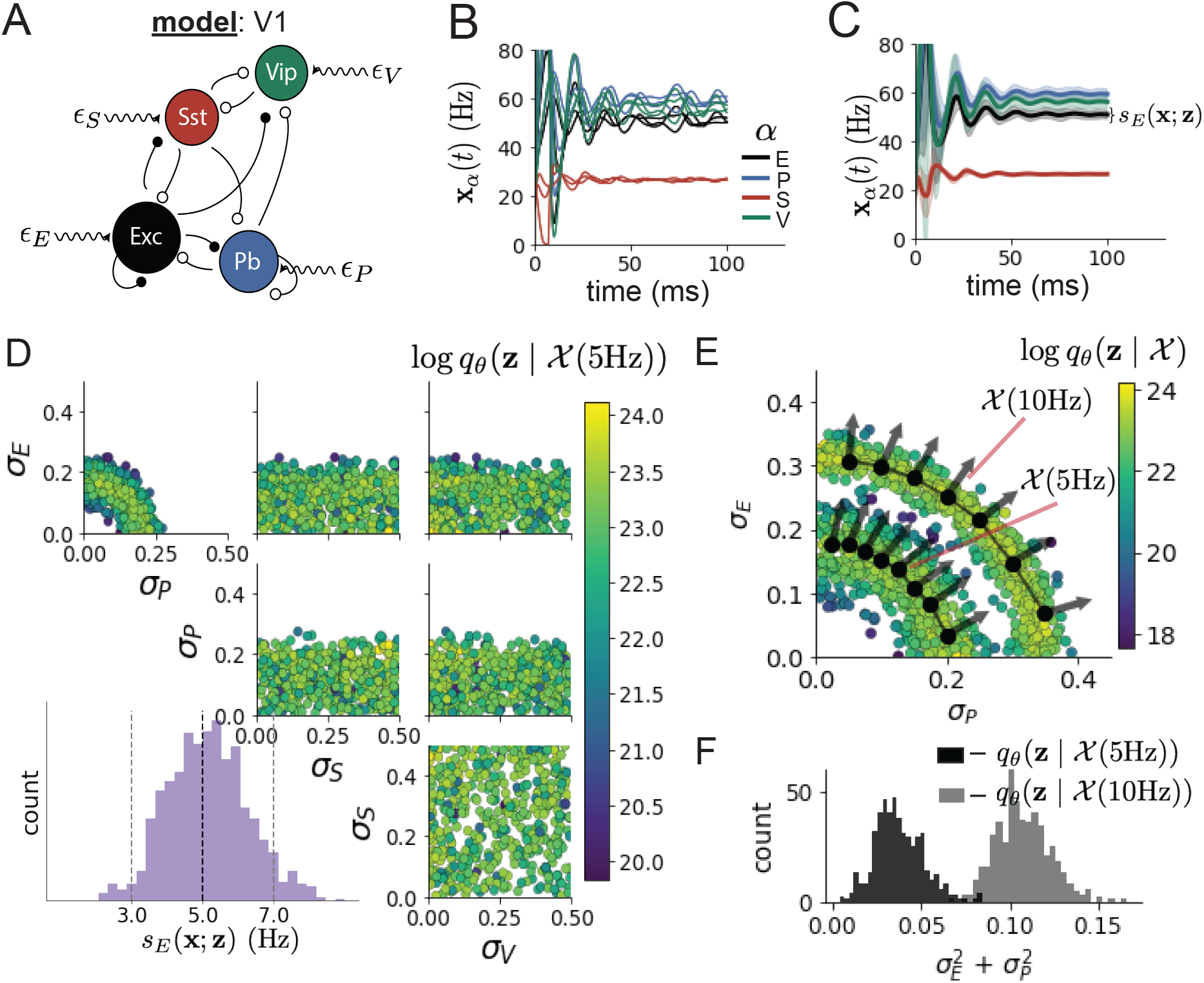
Emergent property inference in the stochastic stabilized supralinear network (SSSN) **A**. Four-population model of primary visual cortex with excitatory (black), parvalbumin (blue), somatostatin (red), and VIP (green) neurons (excitatory and inhibitory projections filled and unfilled, respectively). Some neuron-types largely do not form synaptic projections to others 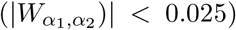. Each neural population receives a baseline input **h**_*b*_, and the E- and P-populations also receive a contrast-dependent input **h**_*c*_. Additionally, each neural population receives a slow noisy input ***ϵ***. **B**. Transient network responses of the SSSN model. Traces are independent trials with varying initialization **x**(0) and noise ***ϵ***. **C**. Mean (solid line) and standard deviation *s_E_*(**x**; **z**) (shading) across 100 trials. **D**. EPI distribution of noise parameters **z** conditioned on E-population variability. The EPI predictive distribution of *s_E_*(**x**; **z**) is show on the bottom-left. **E**. (Top) Enlarged visualization of the *σ_E_*-*σ_P_* marginal distribution of EPI 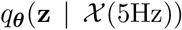 and 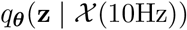. Each black dot shows the mode at each *σ_P_*. The arrows show the most sensitive dimensions of the Hessian evaluated at these modes. **F**. The predictive distributions of 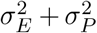 of each inferred distribution 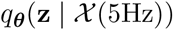 and 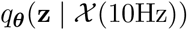.

We considered response variability of a nonlinear dynamical V1 circuit model (Fig. 3A) with a state comprised of each neuron-type population’s rate **x** = [*x_E_, x_P_, x_S_, x_V_*]^⊤^. Each population receives recurrent input *W* **x**, where *W* is the effective connectivity matrix (see Section 5.4) and an external input with mean **h**, which determines population rate via supralinear nonlinearity 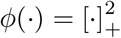. The external input has an additive noisy component ***ϵ*** with variance 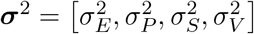. This noise has a slower dynamical timescale *τ*_noise_ > *τ* than the population rate, allowing fluctuations around a stimulus-dependent steady-state (Fig. 3B). This model is the stochastic stabilized supralinear network (SSSN) [59]

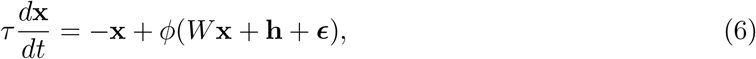

generalized to have multiple inhibitory neuron types. It introduces stochasticity to four neuron-type models of V1 [46]. Stochasticity and inhibitory multiplicity introduce substantial complexity to the mathematical treatment of this problem (see Section 5.4.5) motivating the analysis of this model with EPI. Here, we consider fixed weights *W* and input **h** [47], and study the effect of input variability **z** = [*σ_E_, σ_P_, σ_S_, σ_V_*]^⊤^ on excitatory variability.

We quantify levels of E-population variability by studying two emergent properties

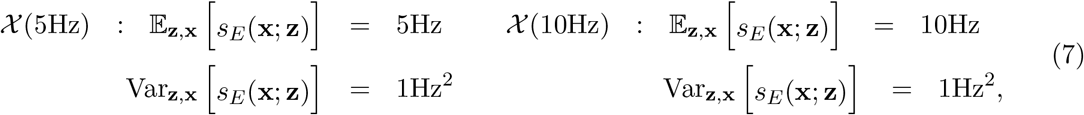

where *s_E_*(**x**; **z**) is the standard deviation of the stochastic *E*-population response about its steady state (Fig. 3C). In the following analyses, we select 1Hz^2^ variance such that the two emergent properties do not overlap in *s_E_*(**z**; **x**).

First, we ran EPI to obtain parameter distribution 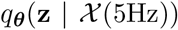 producing E-population variability around 5Hz (Fig. 3D). From the marginal distribution of *σ_E_* and *σ_P_* (Fig. 3D, top-left), we can see that *s_E_*(**x**; **z**) is sensitive to various combinations of *σ_E_* and *σ_P_*. Alternatively, both *σ_S_* and *σ_V_* are degenerate with respect to *s_E_*(**x**; **z**) evidenced by the unexpectedly high variability in those dimensions (Fig. 3D, bottom-right). Together, these observations imply a curved path with respect to *s_E_*(**x**; **z**) of 5Hz, which is indicated by the modes along *σ_P_* (Fig. 3E).

Figure 3E suggests a quadratic relationship in E-population fluctuations and the standard deviation of E- and P-population input; as the square of either *σ_E_* or *σ_P_* increases, the other compensates by decreasing to preserve the level of *s_E_*(**x**; **z**). This quadratic relationship is preserved at greater level of E-population variability 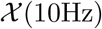 (Fig. 3E and S8). Indeed, the sum of squares of *σ_E_* and *σ_P_* is larger in 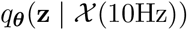 than 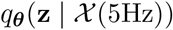 (Fig 3F, *p* < 1 × 10^*−*10^), while the sum of squares of *σ_S_* and *σ_V_* are not significantly different in the two EPI distributions (Fig. S10, *p* = .40), in which parameters were bounded from 0 to 0.5. The strong interaction between E- and P-population input variability on excitatory variability is intriguing, since this circuit exhibits a paradoxical effect in the P-population (and no other inhibitory types) (Fig. S11), meaning that the E-population is P-stabilized. Future research may uncover a link between the population of network stabilization and compensatory interactions governing excitatory variability.

EPI revealed the quadratic dependence of excitatory variability on input variability to the E- and P-populations, as well as its independence to input from the other two inhibitory populations. In a simplified model (*τ* = *τ*_noise_), it can be shown that surfaces of equal variance are ellipsoids as a function of ***σ*** (see Section 5.4.5). Nevertheless, the sensitive and degenerate parameters are intractable to predict mathematically, since the covariance matrix depends on the steady-state solution of the network [59, 66], and terms in the covariance expression increase quadratically with each additional neuron-type population (see also Section 5.4.5). By pointing out this mathematical complexity, we emphasize the value of EPI for gaining understanding about theoretical models when mathematical analysis becomes onerous or impractical.

### 3.5 EPI identifies two regimes of rapid task switching

It has been shown that rats can learn to switch from one behavioral task to the next on randomly interleaved trials [67], and an important question is what neural mechanisms produce this computation. In this experimental setup, rats were given an explicit task cue on each trial, either Pro or Anti. After a delay period, rats were shown a stimulus, and made a context (task) dependent response (Fig. 4A). In the Pro task, rats were required to orient towards the stimulus, while in the Anti task, rats were required to orient away from the stimulus. Pharmacological inactivation of the SC impaired rat performance, and time-specific optogenetic inactivation revealed a crucial role for the SC on the cognitively demanding Anti trials [48]. These results motivated a nonlinear dynamical model of the SC containing four functionally-defined neuron-type populations. In Duan et al. 2019, a computationally intensive procedure was used to obtain a set of 373 connectivity parameters that qualitatively reproduced these optogenetic inactivation results. To build upon the insights of this previous work, we use the probabilistic tools afforded by EPI to identify and characterize two linked, yet distinct regimes of rapid task switching connectivity.

**Figure 4:**
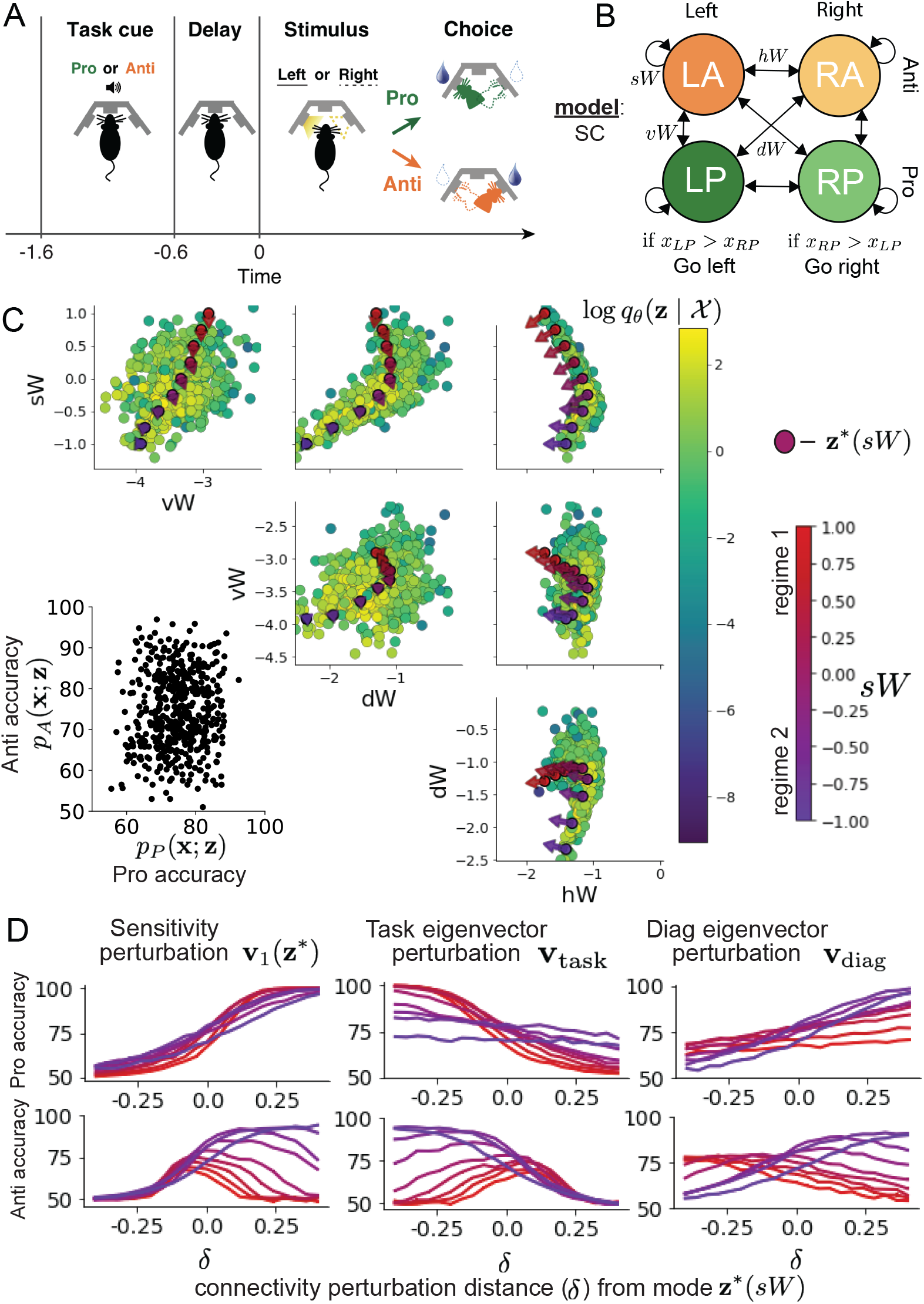
**A**. Rapid task switching behavioral paradigm (see text). **B**. Model of superior colliculus (SC). Neurons: LP - Left Pro, RP - Right Pro, LA - Left Anti, RA - Right Anti. Parameters: *sW* - self, *hW* - horizontal, *vW* -vertical, *dW* - diagonal weights. **C**. The EPI inferred distribution of rapid task switching networks. Red/purple parameters indicate modes **z**\* (*sW*) colored by *sW*. Sensitivity vectors **v**_1_(**z**\*) are shown by arrows. (Bottom-left) EPI predictive distribution of task accuracies. **D**. Mean and standard error (*N*_test_ = 25, bars not visible) of accuracy in Pro (top) and Anti (bottom) tasks after perturbing connectivity away from mode along **v**_1_(**z**\*) (left), **v**_task_ (middle), and **v**_diag_ (right).

In this SC model, there are Pro- and Anti-populations in each hemisphere (left (L) and right (R)) with activity variables **x** = [*x_LP_, x_LA_, x_RP_, x_RA_*]^⊤^ [48]. The connectivity of these populations is parameterized by self *sW*, vertical *vW*, diagonal *dW* and horizontal *hW* connections (Fig. 4B). The input **h** is comprised of a positive cue-dependent signal to the Pro or Anti populations, a positive stimulus-dependent input to either the Left or Right populations, and a choice-period input to the entire network (see Section 5.5.1). Model responses are bounded from 0 to 1 as a function *ϕ* of an internal variable **u**

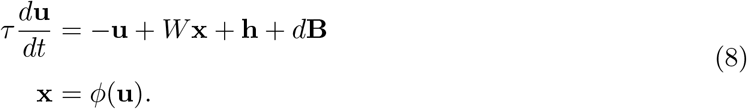

The model responds to the side with greater Pro neuron activation; e.g. the response is left if *x_LP_ > x_RP_* at the end of the trial. Here, we use EPI to determine the network connectivity **z** = [*sW, vW, dW, hW*]^⊤^ that produces rapid task switching.

Rapid task switching is formalized mathematically as an emergent property with two statistics: accuracy in the Pro task *p_P_* (**x**; **z**) and Anti task *p_A_*(**x**; **z**). We stipulate that accuracy be on average .75 in each task with variance .075^2^

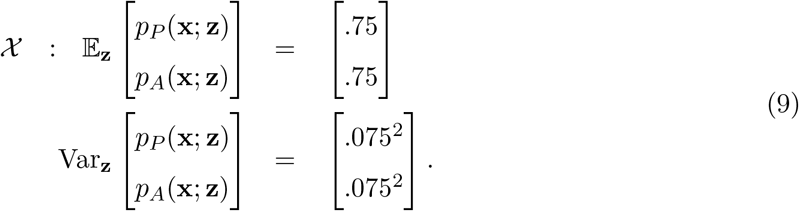

75% accuracy is a realistic level of performance in each task, and with the chosen variance, inferred models will not exhibit fully random responses (50%), nor perfect performance (100%).

The EPI inferred distribution (Fig. 4C) produces Pro and Anti task accuracies (Fig. 4C, bottom-left) consistent with rapid task switching (Equation 9). This parameter distribution has rich structure that is not captured well by simple linear correlations (Fig. S12). Specifically, the shape of the EPI distribution is sharply bent, matching ground truth structure indicated by brute-force sampling (Fig. S18). This is most saliently observed in the marginal distribution of *sW* -*hW* (Fig. 4C top-right), where anticorrelation between *sW* and *hW* switches to correlation with decreasing *sW*. By identifying the modes of the EPI distribution **z**\* (*sW*) at different values of *sW* (Fig. 4C red/purple dots), we can quantify this change in distributional structure with the sensitivity dimension **v**_1_(**z**) (Fig. 4C red/purple arrows). Note that the directionality of these sensitivity dimensions at **z**\* (*sW*) changes distinctly with *sW*, and are perpendicular to the robust dimensions of the EPI distribution that preserve rapid task switching. These two directionalities of sensitivity motivate the distinction of connectivity into two regimes, which produce different types of responses in the Pro and Anti tasks (Fig. S13).

When perturbing connectivity along the sensitivity dimension away from the modes

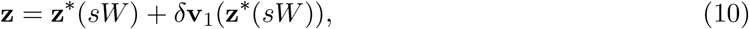

Pro accuracy monotonically increases in both regimes (Fig. 4D, top-left). However, there is a stark difference between regimes in Anti accuracy. Anti accuracy falls in either direction of **v**_1_ in regime 1, yet monotonically increases along with Pro accuracy in regime 2 (Fig. 4D, bottom-left). The sharp change in local structure of the EPI distribution is therefore explained by distinct sensitivities: Anti accuracy diminishes in only one or both directions of the sensitivity perturbation.

To understand the mechanisms differentiating the two regimes, we can make connectivity perturbations along dimensions that only modify a single eigenvalue of the connectivity matrix. These eigenvalues *λ*_all_, *λ*_side_, *λ*_task_, and *λ*_diag_ correspond to connectivity eigenmodes with intuitive roles in processing in this task (Fig. S14A). For example, greater *λ*_task_ will strengthen internal representations of task, while greater *λ*_diag_ will amplify dominance of Pro and Anti pairs in opposite hemispheres (Section 5.5.7). Unlike the sensitivity dimension, the dimensions **v**_*a*_ that perturb isolated connectivity eigenvalues *λ_a_* for *a* ∈ {all, side, task, diag} are independent of **z**\* (*sW*) (see Section 5.5.7), e.g.

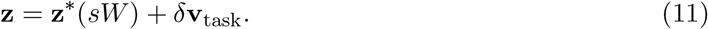

Connectivity perturbation analyses reveal that decreasing *λ*_task_ has a very similar effect on Anti accuracy as perturbations along the sensitivity dimension (Fig. 4D, middle). The similar effects of perturbations along the sensitivity dimension **v**_1_(**z**\*) and reduction of task eigenvalue (via perturbations along −**v**_task_) suggest that there is a carefully tuned strength of task representation in connectivity regime 1, which if disturbed results in random Anti trial responses. Finally, we recognize that increasing *λ*_diag_ has opposite effects on Anti accuracy in each regime (Fig. 4D, right). In the next section, we build on these mechanistic characterizations of each regime by examining their resilience to optogenetic inactivation.

### 3.6 EPI inferred SC connectivities reproduce results from optogenetic inactivation experiments

During the delay period of this task, the circuit must prepare to execute the correct task according to the presented cue. The circuit must then maintain a representation of task throughout the delay period, which is important for correct execution of the Anti task. Duan et al. found that bilateral optogenetic inactivation of SC during the delay period consistently decreased performance in the Anti task, but had no effect on the Pro task (Fig. 5A) [48]. The distribution of connectivities inferred by EPI exhibited this same effect in simulation at high optogenetic strengths *γ*, which reduce the network activities **x**(*t*) by a factor 1 − *γ* (Fig. 5B) (see Section 5.5.8).

**Figure 5:**
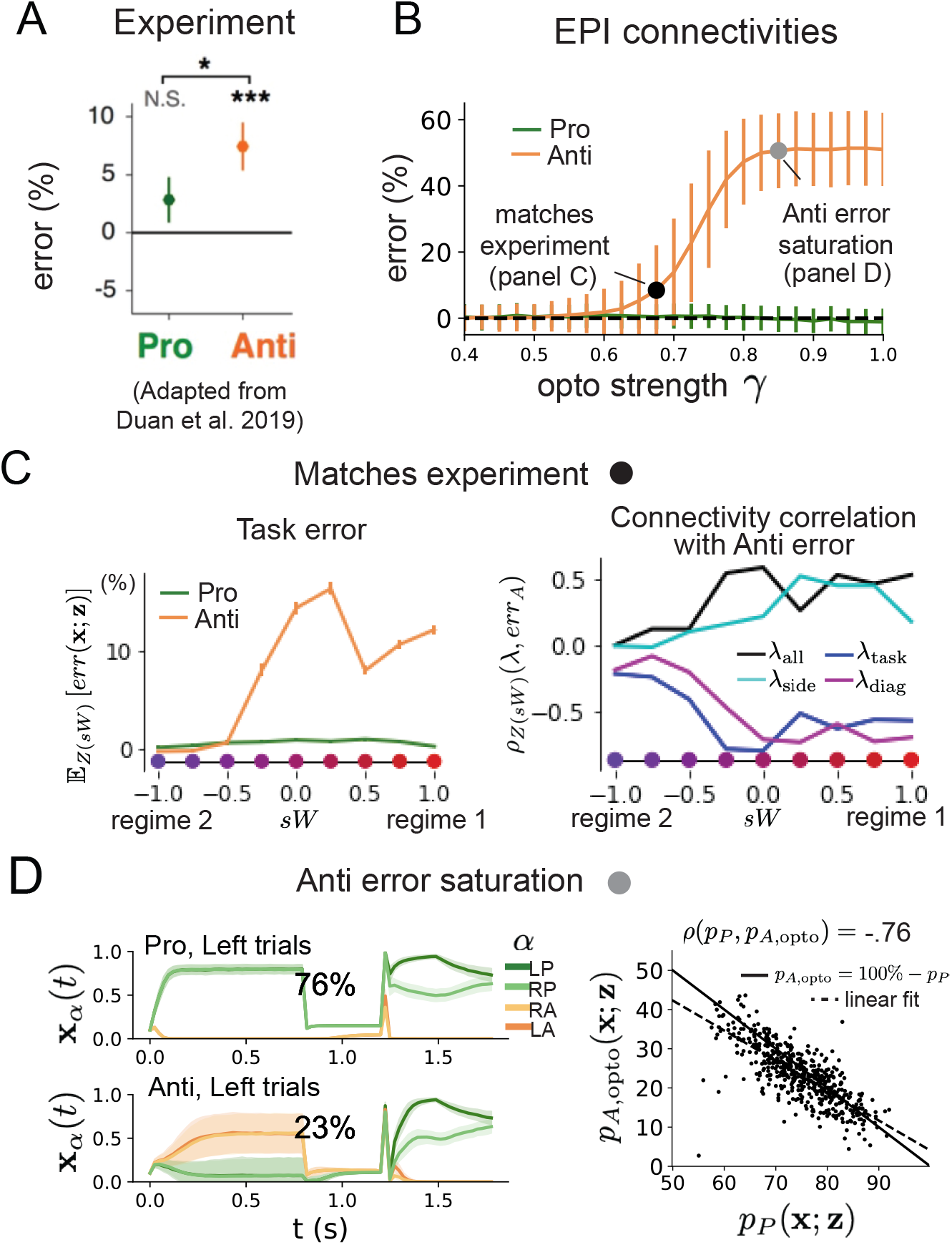
**A**. Mean and standard error (bars) across recording sessions of task error following delay period optogenetic inactivation in rats. **B**. Mean and standard deviation (bars) of task error induced by delay period inactivation of varying optogenetic strength *γ* across the EPI distribution. **C**. (Left) Mean and standard error of Pro and Anti error from regime 1 to regime 2 at *γ* = 0.675. (Right) Correlations of connectivity eigenvalues with Anti error from regime 1 to regime 2 at *γ* = 0.675. **D**. (Left) Mean and standard deviation (shading) of responses of the SC model at the mode of the EPI distribution to delay period inactivation at *γ* = 0.85. Accuracy in Pro (top) and Anti (bottom) task is shown as a percentage. (Right) Anti accuracy following delay period inactivation at *γ* = 0.85 versus accuracy in the Pro task across connectivities in the EPI distribution.

To examine how connectivity affects response to delay period inactivation, we grouped connectivities of the EPI distribution along the continuum linking regimes 1 and 2 of Section 3.5. *Z*(*sW*) is the set of EPI samples for which the closest mode was **z**\* (*sW*) (see Section 5.5.4). In the following analyses, we examine how error, and the influence of connectivity eigenvalue on Anti error change along this continuum of connectivities. Obtaining the parameter samples for these analysis with the learned EPI distribution was more than 20,000 times faster than a brute force approach (see Section 5.5.5).

The mean increase in Anti error of the EPI distribution is closest to the experimentally measured value of 7% at *γ* = 0.675 (Fig. 5B, black dot). At this level of optogenetic strength, regime 1 exhibits an increase in Anti error with delay period silencing (Fig. 5C, left), while regime 2 does not. In regime 1, greater *λ*_task_ and *λ*_diag_ decrease Anti error (Fig. 5C, right). In other words, stronger task representations and diagonal amplification make the SC model more resilient to delay period silencing in the Anti task. This complements the finding from Duan et al. 2019 [48] that *λ*_task_ and *λ*_diag_ improve Anti accuracy.

At roughly *γ* = 0.85 (Fig. 5B, gray dot), the Anti error saturates, while Pro error remains at zero. Following delay period inactivation at this optogenetic strength, there are strong similarities in the responses of Pro and Anti trials during the choice period (Fig. 5D, left). We interpreted these similarities to suggest that delay period inactivation at this saturated level flips the internal representation of task (from Anti to Pro) in the circuit model. A flipped task representation would explain why the Anti error saturates at 50%: the average Anti accuracy in EPI inferred connectivities is 75%, but is 25% when the internal representation is flipped during delay period silencing. This hypothesis prescribes a model of Anti accuracy during delay period silencing of *p*_*A,*opto_ = 100% − *p_P_*, which is fit closely across both regimes of the EPI inferred connectivities (Fig. 5D, right). Similarities between Pro and Anti trial responses were not present at the experiment-matching level of *γ* = 0.675 (Fig. S16 left) and neither was anticorrelation in *p_P_* and *p*_*A,*opto_ (Fig. S16 right).

In summary, the connectivity inferred by EPI to perform rapid task switching replicated results from optogenetic silencing experiments. We found that at levels of optogenetic strength matching experimental levels of Anti error, only one regime actually exhibited the effect. This connectivity regime is less resilient to optogenetic perturbation, and perhaps more biologically realistic. Finally, we characterized the pathology in Anti error that occurs in both regimes when optogenetic strength is increased to high levels, leading to a mechanistic hypothesis that is experimentally testable. The probabilistic tools afforded by EPI yielded this insight: we identified two regimes and the continuum of connectivities between them by taking gradients of parameter probabilities in the EPI distribution, we identified sensitivity dimensions by measuring the Hessian of the EPI distribution, and we obtained many parameter samples at each step along the continuum at an efficient rate.

## 4 Discussion

In neuroscience, machine learning has primarily been used to reveal structure in neural datasets [20]. Careful inference procedures are developed for these statistical models allowing precise, quantitative reasoning, which clarifies the way data informs beliefs about the model parameters. However, these statistical models often lack resemblance to the underlying biology, making it unclear how to go from the structure revealed by these methods, to the neural mechanisms giving rise to it. In contrast, theoretical neuroscience has primarily focused on careful models of neural circuits and the production of emergent properties of computation, rather than measuring structure in neural datasets. In this work, we improve upon parameter inference techniques in theoretical neuroscience with emergent property inference, harnessing deep learning towards parameter inference in neural circuit models (see Section 5.1.1).

Methodology for statistical inference in circuit models has evolved considerably in recent years. Early work used rejection sampling techniques [24–26], but EPI and another recently developed methodology [35] employ deep learning to improve efficiency and provide flexible approximations. SNPE has been used for posterior inference of parameters in circuit models conditioned upon exemplar data used to represent computation, but it does not infer parameter distributions that only produce the computation of interest like EPI (see Section 3.3). When strict control over the predictions of the inferred parameters is necessary, EPI uses a constrained optimization technique [38] (see Section 5.1.4) to make inference conditioned on the emergent property possible.

A key difference between EPI and SNPE, is that EPI uses gradients of the emergent property throughout optimization. In Section 3.3, we showed that such gradients confer beneficial scaling properties, but a concern remains that emergent property gradients may be too computationally intensive. Even in a case of close biophysical realism with an expensive emergent property gradient, EPI was run successfully on intermediate hub frequency in a 5-neuron subcircuit model of the STG (Section 3.1). However, conditioning on the pyloric rhythm [68] in a model of the pyloric subnetwork model [15] proved to be prohibitive with EPI. The pyloric subnetwork requires many time steps for simulation and many key emergent property statistics (e.g. burst duration and phase gap) are not calculable or easily approximated with differentiable functions. In such cases, SNPE, which does not require differentiability of the emergent property, has proven useful [35]. In summary, choice of deep inference technique should consider emergent property complexity and differentiability, dimensionality of parameter space, and the importance of constraining the model behavior predicted by the inferred parameter distribution.

In this paper, we demonstrate the value of deep inference for parameter sensitivity analyses at both the local and global level. With these techniques, flexible deep probability distributions are optimized to capture global structure by approximating the full distribution of suitable parameters. Importantly, the local structure of this deep probability distribution can be quantified at any parameter choice, offering instant sensitivity measurements after fitting. For example, the global structure captured by EPI revealed two distinct parameter regimes, which had different local structure quantified by the deep probability distribution (see Section 5.5). In comparison, bayesian MCMC is considered a popular approach for capturing global parameter structure [69], but there is no variational approximation (the deep probability distribution in EPI), so sensitivity information is not queryable and sampling remains slow after convergence. Local sensitivity analyses (e.g. [27]) may be performed independently at individual parameter samples, but these methods alone do not capture the full picture in nonlinear, complex distributions. In contrast, deep inference yields a probability distribution that produces a wholistic assessment of parameter sensitivity at the local and global level, which we used in this study to make novel insights into a range of theoretical models. Together, the abilities to condition upon emergent properties, the efficient inference algorithm, and the capacity for parameter sensitivity analyses make EPI a useful method for addressing inverse problems in theoretical neuroscience.

## Acknowledgements

This work was funded by NSF Graduate Research Fellowship, DGE-1644869, McKnight Endowment Fund, NIH NINDS 5R01NS100066, Simons Foundation 542963, NSF NeuroNex Award, DBI-1707398, The Gatsby Charitable Foundation, Simons Collaboration on the Global Brain Postdoctoral Fellowship, Chinese Postdoctoral Science Foundation, and International Exchange Program Fellowship. We also acknowledge the Marine Biological Laboratory Methods in Computational Neuroscience Course, where this work was discussed and explored in its early stages. Helpful conversations were had with Larry Abbott, Stephen Baccus, James Fitzgerald, Gabrielle Gutierrez, Francesca Mastrogiuseppe, Srdjan Ostojic, Liam Paninski, and Dhruva Raman.

## Data availability statement

The datasets generated during and/or analyzed during the current study are available from the corresponding author upon reasonable request.

## Code availability statement

All software written for the current study is available at https://github.com/cunningham-lab/epi.

## 5 Methods

### 5.1 Emergent property inference (EPI)

Solving inverse problems is an important part of theoretical neuroscience, since we must understand how neural circuit models and their parameter choices produce computations. Recently, research on machine learning methodology for neuroscience has focused on finding latent structure in large-scale neural datasets, while research in theoretical neuroscience generally focuses on developing precise neural circuit models that can produce computations of interest. By quantifying computation into an *emergent property* through statistics of the emergent activity of neural circuit models, we can adapt the modern technique of deep probabilistic inference towards solving inverse problems in theoretical neuroscience. Here, we introduce a novel method for statistical inference, which uses deep networks to learn parameter distributions constrained to produce emergent properties of computation.

Consider model parameterization **z**, which is a collection of scientifically meaningful variables that govern the complex simulation of data **x**. For example (see Section 3.1), **z** may be the electrical conductance parameters of an STG subcircuit, and **x** the evolving membrane potentials of the five neurons. In terms of statistical modeling, this circuit model has an intractable likelihood *p*(**x** | **z**), which is predicated by the stochastic differential equations that define the model. From a theoretical perspective, we are less concerned about the likelihood of an exemplar dataset **x**, but rather the emergent property of intermediate hub frequency (which implies a consistent dataset **x**).

In this work, emergent properties 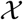 are defined through the choice of emergent property statistic *f* (**x**; **z**) (which is a vector of one or more statistics), and its means ***μ***, and variances ***σ***^2^:

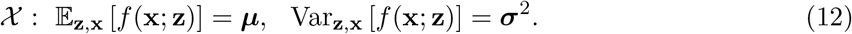

In general, an emergent property may be a collection of first-, second-, or higher-order moments of a group of statistics, but this study focuses on the case written in Equation 12. In the STG example, intermediate hub frequency is defined by mean and variance constraints on the statistic of hub neuron frequency *ω*_hub_(**x**; **z**) (Equations 2 and 3). Precisely, the emergent property statistics *f* (**x**; **z**) must have means ***μ*** and variances ***σ***^2^ over the EPI distribution of parameters (**z** ~ *q_**θ**_*(**z**)) and the data produced by those parameters (**x** ~ *p*(**x** | **z**)), where the inferred parameter distribution *q_**θ**_*(**z**) itself is parameterized by deep network weights and biases ***θ***.

In EPI, a deep probability distribution *q_**θ**_*(**z**) is optimized to approximate the parameter distribution producing the emergent property 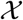. In contrast to simpler classes of distributions like the gaussian or mixture of gaussians, deep probability distributions are far more flexible and capable of fitting rich structure [36, 37]. In deep probability distributions, a simple random variable **z**_0_ ~ *q*_0_(**z**_0_) (we choose an isotropic gaussian) is mapped deterministically via a sequence of deep neural network layers (*g*_1_, .. *g_l_*) parameterized by weights and biases ***θ*** to the support of the distribution of interest:

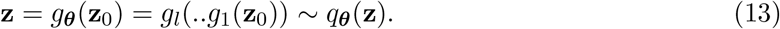

Such deep probability distributions embed the inferred distribution in a deep network. Once optimized, this deep network representation of a distribution has remarkably useful properties: fast sampling and probability evaluations. Importantly, fast probability evaluations confer fast gradient and Hessian calculations as well.

Given this choice of circuit model and emergent property 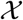, *q_**θ**_*(**z**) is optimized via the neural network parameters ***θ*** to find a maximally entropic distribution 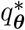 within the deep variational family 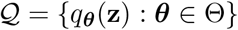 that produces the emergent property 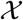:

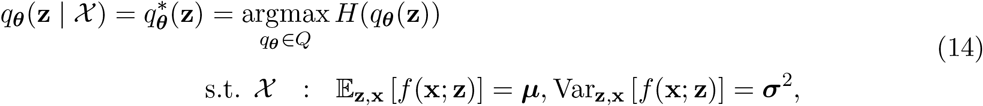

where 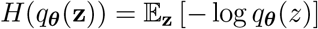 is entropy. By maximizing the entropy of the inferred distribution *q_**θ**_*, we select the most random distribution in family 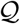 that satisfies the constraints of the emergent property. Since entropy is maximized in Equation 14, EPI is equivalent to bayesian variational inference (see Section 5.1.6), which is why we specify the inferred distribution of EPI as conditioned upon emergent property 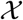 with the notation 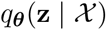. To run this constrained optimization, we use an augmented lagrangian objective, which is the standard approach for constrained optimization [70], and the approach taken to fit Maximum Entropy Flow Networks (MEFNs) [38]. This procedure is detailed in Section 5.1.4 and the pseudocode in Algorithm 1.

In the remainder of Section 5.1, we will explain the finer details and motivation of the EPI method. First, we explain related approaches and what EPI introduces to this domain (Section 5.1.1). Second, we describe the special class of deep probability distributions used in EPI called normalizing flows (Section 5.1.2). Then, we establish the known relationship between maximum entropy distributions and exponential families (Section 5.1.3). Next, we explain the constrained optimization technique used to solve Equation 14 (Section 5.1.4). Then, we demonstrate the details of this optimization in a toy example (Section 5.1.5). Finally, we explain how EPI is equivalent to variational inference (Section 5.1.6).

#### 5.1.1 Related approaches

When bayesian inference problems lack conjugacy, scientists use approximate inference methods like variational inference (VI) [71] and Markov chain Monte Carlo (MCMC) [72,73]. After optimization, variational methods return a parameterized posterior distribution, which we can analyze. Also, the variational approximation is often chosen such that it permits fast sampling. In contrast MCMC methods only produce samples from the approximated posterior distribution. No parameterized distribution is estimated, and additional samples are always generated with the same sampling complexity. Inference in models defined by systems of differential has been demonstrated with MCMC [69], although this approach requires tractable likelihoods. Advancements have introduced sampling [74], likelihood approximation [75], and uncertainty quantification techniques [76] to make MCMC approaches more efficient and expand the class of applicable models.

Simulation-based inference [56] is model parameter inference in the absence of a tractable likelihood function. The most prevalent approach to simulation-based inference is approximate bayesian computation (ABC) [24], in which satisfactory parameter samples are kept from random prior sampling according to a rejection heuristic. The obtained set of parameters do not have a probabilities, and further insight about the model must be gained from examination of the parameter set and their generated activity. Methodological advances to ABC methods have come through the use of Markov chain Monte Carlo (MCMC-ABC) [25] and sequential Monte Carlo (SMC-ABC) [26] sampling techniques. SMC-ABC is considered state-of-the-art ABC, yet this approach still struggles to scale in dimensionality [55] (cf. Fig. 2). Still, this method has enjoyed much success in systems biology [77]. Furthermore, once a parameter set has been obtained by SMC-ABC from a finite set of particles, the SMC-ABC algorithm must be run again from scratch with a new population of initialized particles to obtain additional samples.

For scientific model analysis, we seek a parameter distribution represented by an approximating distribution as in variational inference [71]: a variational approximation that once optimized yields fast analytic calculations and samples. For the reasons described above, ABC and MCMC techniques are not suitable, since they only produce a set of parameter samples lacking probabilities and have unchanging sampling rate. EPI infers parameters in circuit models using the MEFN [38] algorithm with a deep variational approximation. The deep neural network of EPI (Fig. 1E) defines the parametric form (with weights and biases as variational parameters ***θ***) of the variational approximation of the inferred parameter distribution *q_**θ**_*(**z** | **x**). The EPI optimization is enabled using stochastic gradient techniques in the spirit of likelihood-free variational inference [34]. The analytic relationship between EPI and variational inference is explained in Section 5.1.6.

We note that, during our preparation and early presentation of this work [78, 79], another work has arisen with broadly similar goals: bringing statistical inference to mechanistic models of neural circuits [35, 80, 81]. We are encouraged by this general problem being recognized by others in the community, and we emphasize that these works offer complementary neuroscientific contributions (different theoretical models of focus) and use different technical methodologies (ours is built on our prior work [38], theirs similarly [82]).

The method EPI differs from SNPE in some key ways. SNPE belongs to a “sequential” class of recently developed simulation-based inference methods in which two neural networks are used for posterior inference. This first neural network is a deep probability distribution (normalizing flow) used to estimate the posterior *p*(**z** | **x**) (SNPE) or the likelihood *p*(**x** | **z**) (sequential neural likelihood (SNL) [83]). A recent approach uses an unconstrained neural network to estimate the likelihood ratio (sequential neural ratio estimation (SNRE) [84]). In SNL and SNRE, MCMC sampling techniques are used to obtain samples from the approximated posterior. This contrasts with EPI and SNPE, which use deep probability distributions to model parameters, which facilitates immediate measurements of sample probability, gradient, or Hessian for system analysis. The second neural network in this sequential class of methods is the amortizer. This unconstrained deep network maps data **x**(or statistics *f* (**x**; **z**) or model parameters **z**) to the weights and biases of the first neural network. These methods are optimized on a conditional density (or ratio) estimation objective. The data used to optimize this objective are generated via an adaptive procedure, in which training data pairs (**x**_*i*_, **z**_*i*_) become sequentially closer to the true data and posterior.

The approximating fidelity of the deep probability distribution in sequential approaches is optimized to generalize across the training distribution of the conditioning variable. This generalization property of the sequential methods can reduce the accuracy at the singular posterior of interest. Whereas in EPI, the entire expressivity of the deep probability distribution is dedicated to learning a single distribution as well as possible. The well-known inverse mapping problem of exponential families [85] prohibits an amortization-based approach in EPI, since EPI learns an exponential family distribution parameterized by its mean (in contrast to its natural parameter, see Section 5.1.3). However, we have shown that the same two-network architecture of the sequential simulation-based inference methods can be used for amortized inference in intractable exponential family posteriors when using their natural parameterization [86].

Finally, one important differentiating factor between EPI and sequential simulation-based inference methods is that EPI leverages gradients ∇_**z**_*f* (**x**; **z**) during optimization. These gradients can improve convergence time and scalability, as we have shown on an example conditioning low-rank RNN connectivity on the property of stable amplification (see Section 3.3). With EPI, we prove out the suggestion that a deep inference technique can improve efficiency by leveraging these emergent property gradients when they are tractable. Sequential simulation-based inference techniques may be better suited for scientific problems where ∇_**z**_*f* (**x**; **z**) is intractable or unavailable, like when there is a nondifferentiable emergent property. However, the sequential simulation-based inference techniques cannot constrain the predictions of the inferred distribution in the manner of EPI.

Structural identifiability analysis involves the measurement of sensitivity and unidentifiabilities in scientific models. Around a single parameter choice, one can measure the Jacobian. One approach for this calculation that scales well is EAR [28]. A popular efficient approach for systems of ODEs has been neural ODE adjoint [87] and its stochastic adaptation [88]. Casting identifiability as a statistical estimation problem, the profile likelihood works via iterated optimization while holding parameters fixed [27]. An exciting recent method is capable of recovering the functional form of such unidentifiabilities away from a point by following degenerate dimensions of the fisher information matrix [30]. Global structural non-identifiabilities can be found for models with polynomial or rational dynamics equations using DAISY [89], or through mean optimal transformations [90]. With EPI, we have all the benefits given by a statistical inference method plus the ability to query the first- or second-order gradient of the probability of the inferred distribution at any chosen parameter value. The second-order gradient of the log probability (the Hessian), which is directly afforded by EPI distributions, produces quantified information about parametric sensitivity of the emergent property in parameter space (see Section 3.2).

#### 5.1.2 Deep probability distributions and normalizing flows

Deep probability distributions are comprised of multiple layers of fully connected neural networks (Equation 13). When each neural network layer is restricted to be a bijective function, the sample density can be calculated using the change of variables formula at each layer of the network. For **z**_*i*_ = *g_i_*(**z**_**i***−***1**_),

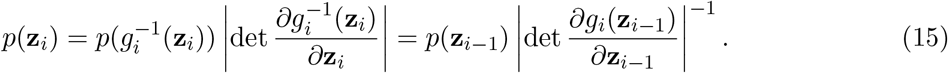

However, this computation has cubic complexity in dimensionality for fully connected layers. By restricting our layers to normalizing flows [36, 37] – bijective functions with fast log determinant Jacobian computations, which confer a fast calculation of the sample log probability. Fast log probability calculation confers efficient optimization of the maximum entropy objective (see Section 5.1.4).

We use the real NVP [39] normalizing flow class, because its coupling architecture confers both fast sampling (forward) and fast log probability evaluation (backward). Fast probability evaluation facilitates fast gradient and Hessian evaluation of log probability throughout parameter space. Glow permutations were used in between coupling stages [40]. This is in contrast to autoregressive architectures [91, 92], in which only one of the forward or backward passes can be efficient. In this work, normalizing flows are used as flexible parameter distribution approximations *q_**θ**_*(**z**) having weights and biases ***θ***. We specify the architecture used in each application by the number of real NVP affine coupling stages, and the number of neural network layers and units per layer of the conditioning functions.

When calculating Hessians of log probabilities in deep probability distributions, it is important to consider the normalizing flow architecture. With autoregressive architectures [91, 92], fast sampling and fast log probability evaluations are mutually exclusive. That makes these architectures undesirable for EPI, where efficient sampling is important for optimization, and log probability evaluation speed predicates the efficiency of gradient and Hessian calculations. With real NVP coupling architectures, we get both fast sampling and fast Hessians making both optimization and scientific analysis efficient.

#### 5.1.3 Maximum entropy distributions and exponential families

The inferred distribution of EPI is a maximum entropy distribution, which have fundamental links to exponential family distributions. A maximum entropy distribution of form:

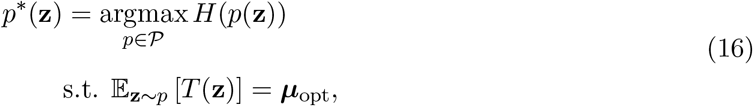

where *T* (**z**) is the sufficient statistics vector and ***μ***_opt_ a vector of their mean values, will have probability density in the exponential family:

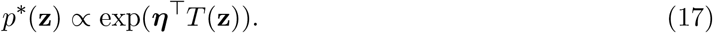

The mappings between the mean parameterization ***μ***_opt_ and the natural parameterization ***η*** are formally hard to identify except in special cases [85].

In this manuscript, emergent properties are defined by statistics *f* (**x**; **z**) having a fixed mean ***μ*** and variance ***σ***^2^ as in Equation 12. The variance constraint is a second moment constraint on *f* (**x**; **z**):

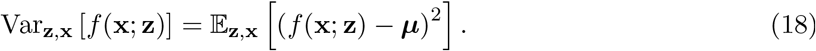

As a general maximum entropy distribution (Equation 16), the sufficient statistics vector contains both first and second order moments of *f* (**x**; **z**)

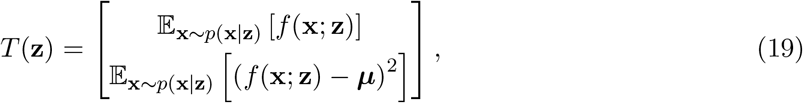

which are constrained to the chosen means and variances

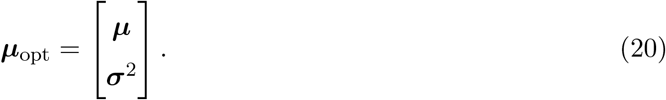

Thus, ***μ***_opt_ is used to denote the mean parameter of the maximum entropy distribution defined by the emergent property (all constraints), while ***μ*** is only the mean of *f* (**x**; **z**). The subscript “opt” of ***μ***_opt_ is chosen since it contains all of the constraint values to which the EPI optimization algorithm must adhere.

#### 5.1.4 Augmented lagrangian optimization

To optimize *q_**θ**_*(**z**) in Equation 14, the constrained maximum entropy optimization is executed using the augmented lagrangian method. The following objective is minimized:

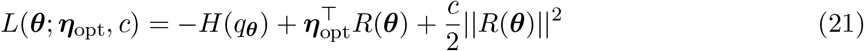

where there are average constraint violations

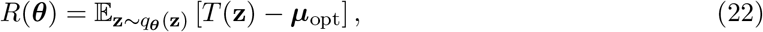

***η***_opt_ ∈ R^*m*^ are the lagrange multipliers where *m* is the number of total constraints

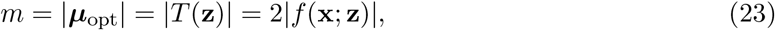

and *c* is the penalty coefficient. The mean parameter ***μ***_opt_ and sufficient statistics *T* (**z**) are determined by the means ***μ*** and variances ***σ***^2^ of the emergent property statistics *f* (**x**; **z**) defined in Equation 14. Specifically, *T* (**z**) is a concatenation of the first and second moments (Equation 19) and ***μ***_opt_ is a concatenation of their constraints ***μ*** and ***σ***^2^ (Equation 20). (Although, note that this algorithm is written for general *T* (**z**) and ***μ***_opt_ to satisfy the more general class of emergent properties.) The lagrange multipliers ***η***_opt_ are closely related to the natural parameters ***η*** of exponential families (see Section 5.1.6). Weights and biases ***θ*** of the deep probability distribution are optimized according to Equation 21 using the Adam optimizer with learning rate 10^*−*3^ [93].

The gradient with respect to entropy *H*(*q_**θ**_*(**z**)) can be expressed using the reparameterization trick as an expectation of the negative log density of parameter samples **z** over the randomness in the parameterless initial distribution *q*_0_(**z**_0_):

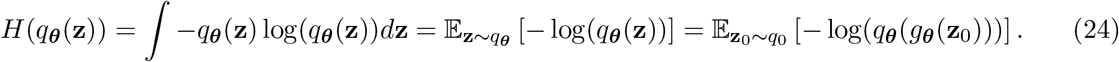

Thus, the gradient of the entropy of the deep probability distribution can be estimated as an average of gradients with respect to the base distribution **z**_0_:

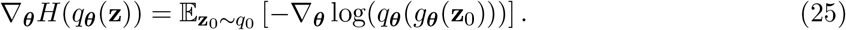

The gradients of the log density of the deep probability distribution are tractable through the use of normalizing flows (see Section 5.1.2).

The full EPI optimization algorithm is detailed in Algorithm 1. The lagrangian parameters ***η***_opt_ are initialized to zero and adapted following each augmented lagrangian epoch, which is a period of optimization with fixed (***η***_opt_, *c*) for a given number of stochastic gradient descent (SGD) iterations. A low value of *c* is used initially, and conditionally increased after each epoch based on constraint error reduction. The penalty coefficient is updated based on the result of a hypothesis test regarding the reduction in constraint violation. The p-value of 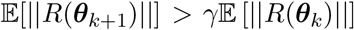 is computed, and *c_k_*_+1_ is updated to *βc_k_* with probability 1 − *p*. The other update rule is 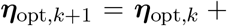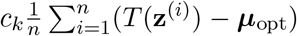 given a batch size *n* and **z**^(*i*)^ ~ *q_**θ**_*(**z**). Throughout the study, *γ* = 0.25, while *β* was chosen to be either 2 or 4. The batch size of EPI also varied according to application.

In general, *c* and ***η***_opt_ should start at values encouraging entropic growth early in optimization. With each training epoch in which the update rule for *c* is invoked, the constraint satisfaction terms are increasingly weighted, which generally results in decreased entropy (e.g. see Figure S1C). This encourages the discovery of suitable regions of parameter space, and the subsequent refinement of the distribution to produce the emergent property. The momentum parameters of the Adam optimizer are reset at the end of each augmented lagrangian epoch, which proceeds for *i*_max_ iterations. In this work, we used a maximum number of augmented lagrangian epochs *k*_max_ >= 5.

Rather than starting optimization from some ***θ*** drawn from a randomized distribution, we found that initializing *q_**θ**_*(**z**) to approximate an isotropic gaussian distribution conferred more stable, consistent optimization. The parameters of the gaussian initialization were chosen on an application-specific basis. Throughout the study, we chose isotropic Gaussian initializations with mean ***μ***_init_ at the center of the support of the distribution and some variance 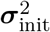, except for one case, where an initialization informed by random search was used (see Section 5.2). Deep probability distributions were fit to these gaussian initializations using 10,000 iterations of stochastic gradient descent on the evidence lower bound (as in [86]) with Adam optimizer and a learning rate of 10^*−*3^.

**Algorithm 1:**
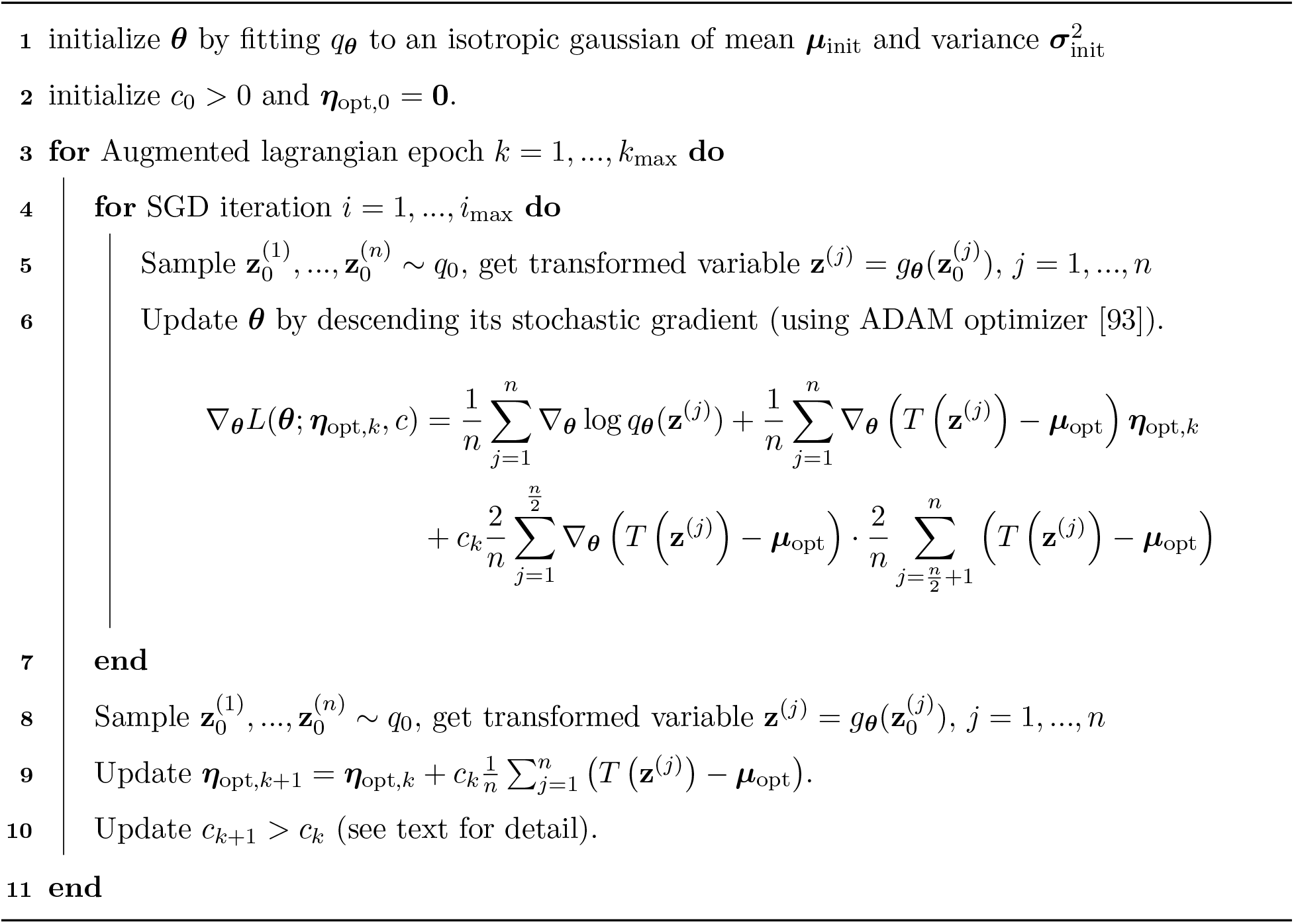
Emergent property inference

To assess whether the EPI distribution *q_**θ**_*(**z**) produces the emergent property, we assess whether each individual constraint on the means and variances of *f* (**x**; **z**) is satisfied. We consider the EPI to have converged when a null hypothesis test of constraint violations *R*(***θ***)_*i*_ being zero is accepted for all constraints *i* ∈ {1, *…, m*} at a significance threshold *α* = 0.05. This significance threshold is adjusted through Bonferroni correction according to the number of constraints *m*. The p-values for each constraint are calculated according to a two-tailed nonparametric test, where 200 estimations of the sample mean *R*(***θ***)^*i*^ are made using *N*_test_ samples of **z** ~ *q_**θ**_*(**z**) at the end of the augmented lagrangian epoch. Of all *k*_max_ augmented lagrangian epochs, we select the EPI inferred distribution as that which satisfies the convergence criteria and has greatest entropy.

When assessing the suitability of EPI for a particular modeling question, there are some important technical considerations. First and foremost, as in any optimization problem, the defined emergent property should always be appropriately conditioned (constraints should not have wildly different units). Furthermore, if the program is underconstrained (not enough constraints), the distribution grows (in entropy) unstably unless mapped to a finite support. If overconstrained, there is no parameter set producing the emergent property, and EPI optimization will fail (appropriately).

#### 5.1.5 Example: 2D LDS

To gain intuition for EPI, consider a two-dimensional linear dynamical system (2D LDS) model (Fig. S1A):

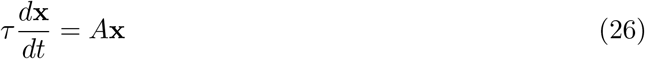

with

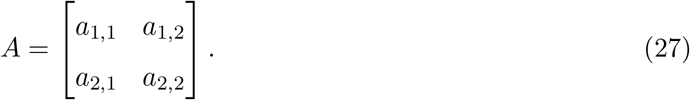

To run EPI with the dynamics matrix elements as the free parameters **z** = [*a*_1,1_, *a*_1,2_, *a*_2,1_, *a*_2,2_] (fixing *τ* = 1s), the emergent property statistics *f* (**x**; **z**) were chosen to contain parts of the primary eigenvalue of *A*, which predicate frequency, imag(*λ*_1_), and the growth/decay, real(*λ*_1_), of the system

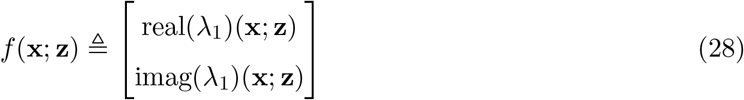

*λ*_1_ is the eigenvalue of greatest real part when the imaginary component is zero, and alternatively that of positive imaginary component when the eigenvalues are complex conjugate pairs. To learn the distribution of real entries of *A* that produce a band of oscillating systems around 1Hz, we formalized this emergent property as real(*λ*_1_) having mean zero with variance 0.25^2^, and the oscillation frequency 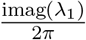 having mean 1Hz with variance 0.1Hz^2^:

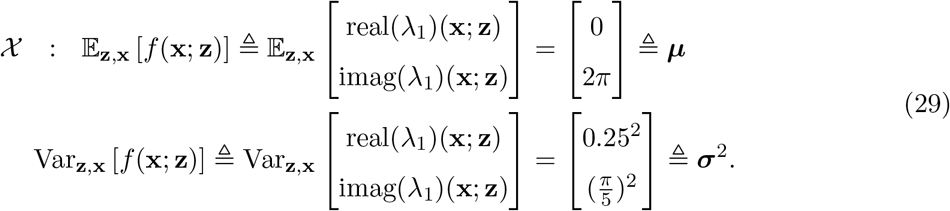

To write the emergent property 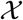 in the form required for the augmented lagrangian optimization (Section 5.1.4), we concatenate these first and second moment constraints into a vector of sufficient statistics *T* (**z**) and constraint values ***μ***_opt_.

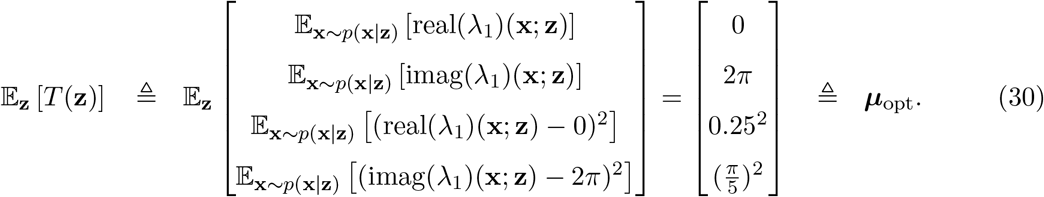

From now on in all scientific applications (Sections 5.2–5.5, we specify how the EPI optimization was setup by specifying *f* (**x**; **z**), ***μ***, and ***σ***^2^.

Unlike the models we presented in the main text, this model admits an analytical form for the mean emergent property statistics given parameter **z**, since the eigenvalues can be calculated using the quadratic formula:

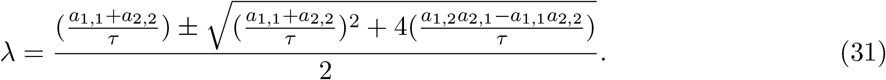

We study this example, because the inferred distribution is curved and multimodal, and we can compare the result of EPI to analytically derived contours of the emergent property statistics.

Despite the simple analytic form of the emergent property statistics, the EPI distribution in this example is not simply determined. Although 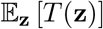 is calculable directly via a closed form function, the distribution 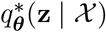 cannot be derived directly. This fact is due to the formally hard problem of the backward mapping: finding the natural parameters ***η*** from the mean parameters ***μ*** of an exponential family distribution [85]. Instead, we used EPI to approximate this distribution (Fig. S1B). We used a real NVP normalizing flow architecture three coupling layers and two-layer neural networks of 50 units per layer, mapped onto a support of *z_i_* ∈ [−10, 10]. (see Section 5.1.2).

**Figure S1:**
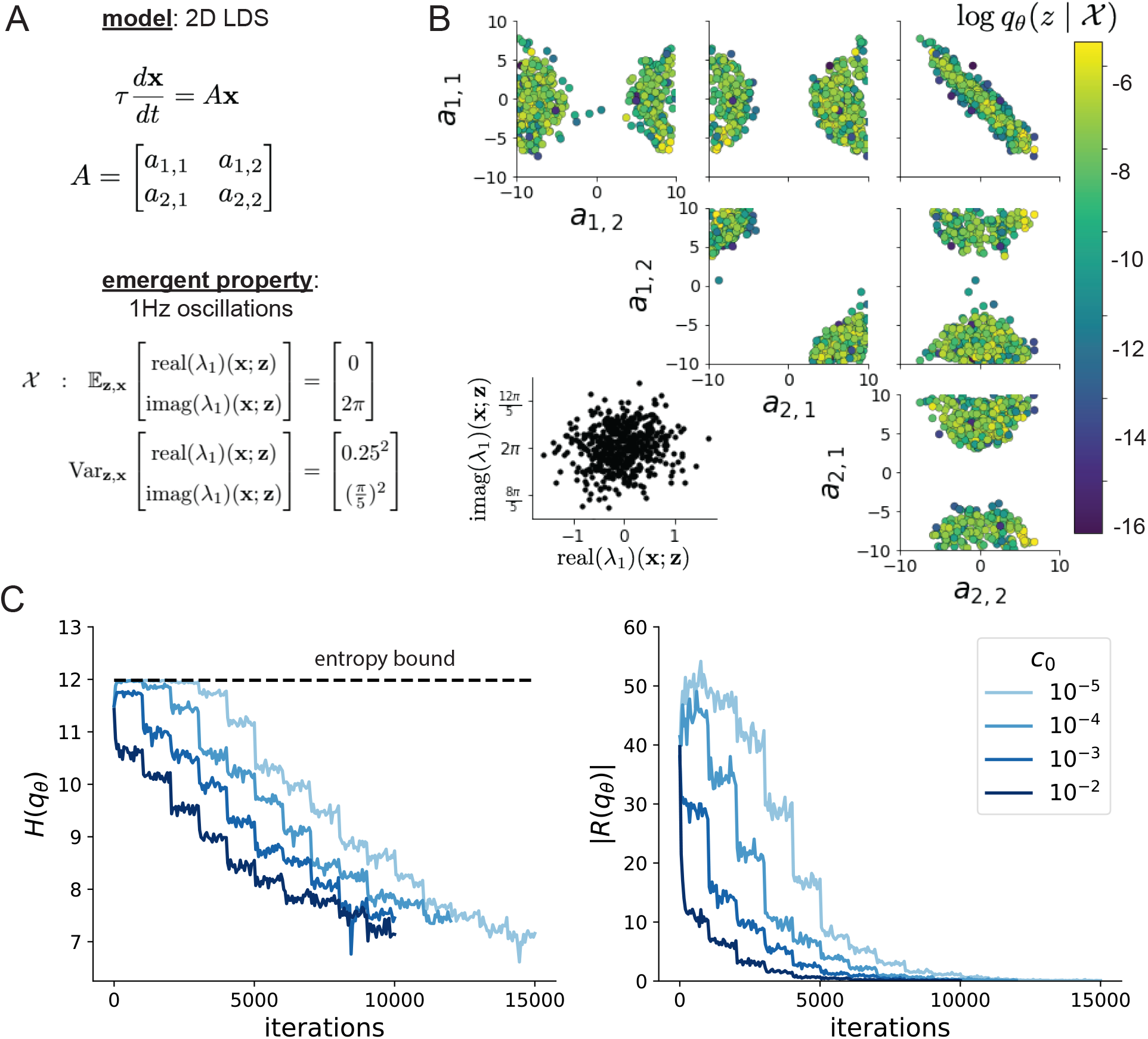
**A**. Two-dimensional linear dynamical system model, where real entries of the dynamics matrix *A* are the parameters. **B**. The EPI distribution for a two-dimensional linear dynamical system with *τ* = 1 that produces an average of 1Hz oscillations with some small amount of variance. Dashed lines indicate the parameter axes. **C**. Entropy throughout the optimization. At the beginning of each augmented lagrangian epoch (*i*_max_ = 2, 000 iterations), the entropy dipped due to the shifted optimization manifold where emergent property constraint satisfaction is increasingly weighted. **D**. Emergent property moments throughout optimization. At the beginning of each augmented lagrangian epoch, the emergent property moments adjust closer to their constraints.

Even this relatively simple system has nontrivial (though intuitively sensible) structure in the parameter distribution. To validate our method, we analytically derived the contours of the probability density from the emergent property statistics and values. In the *a*_1,1_-*a*_2,2_ plane, the black line at 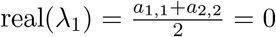, dashed black line at the standard deviation 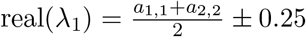, and the dashed gray line at twice the standard deviation 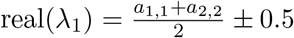 follow the contour of probability density of the samples (Fig. S2A). The distribution precisely reflects the desired statistical constraints and model degeneracy in the sum of *a*_1,1_ and *a*_2,2_. Intuitively, the parameters equivalent with respect to emergent property statistic real(*λ*_1_) have similar log densities.

To explain the bimodality of the EPI distribution, we examined the imaginary component of *λ*_1_. When real(*λ*_1_) = *a*_1,1_ + *a*_2,2_ = 0 (which is the case on average in 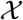), we have

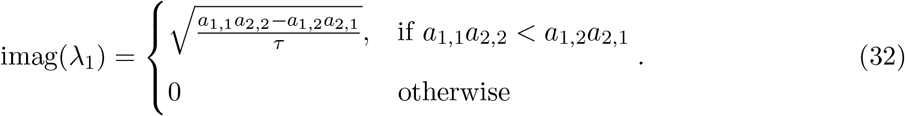

In Figure S2B, we plot the contours of imag(*λ*_1_) where *a*_1,1_*a*_2,2_ is fixed to 0 at one standard deviation (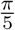, black dashed) and two standard deviations (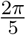, gray dashed) from the mean of 2*π*. This validates the curved multimodal structure of the inferred distribution learned through EPI. Subtler combinations of model and emergent property will have more complexity, further motivating the use of EPI for understanding these systems. As we expect, the distribution results in samples of two-dimensional linear systems oscillating near 1Hz (Fig. S3).

#### 5.1.6 EPI as variational inference

In variational inference, a posterior approximation 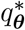 is chosen from within some variational family 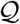 to be as close as possible to the posterior under the KL divergence criteria

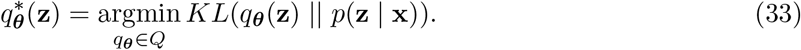

This KL divergence can be written in terms of entropy of the variational approximation:

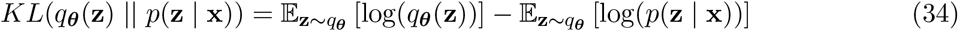

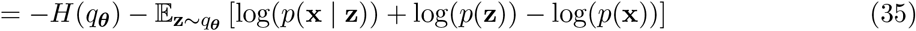

**Figure S2:**
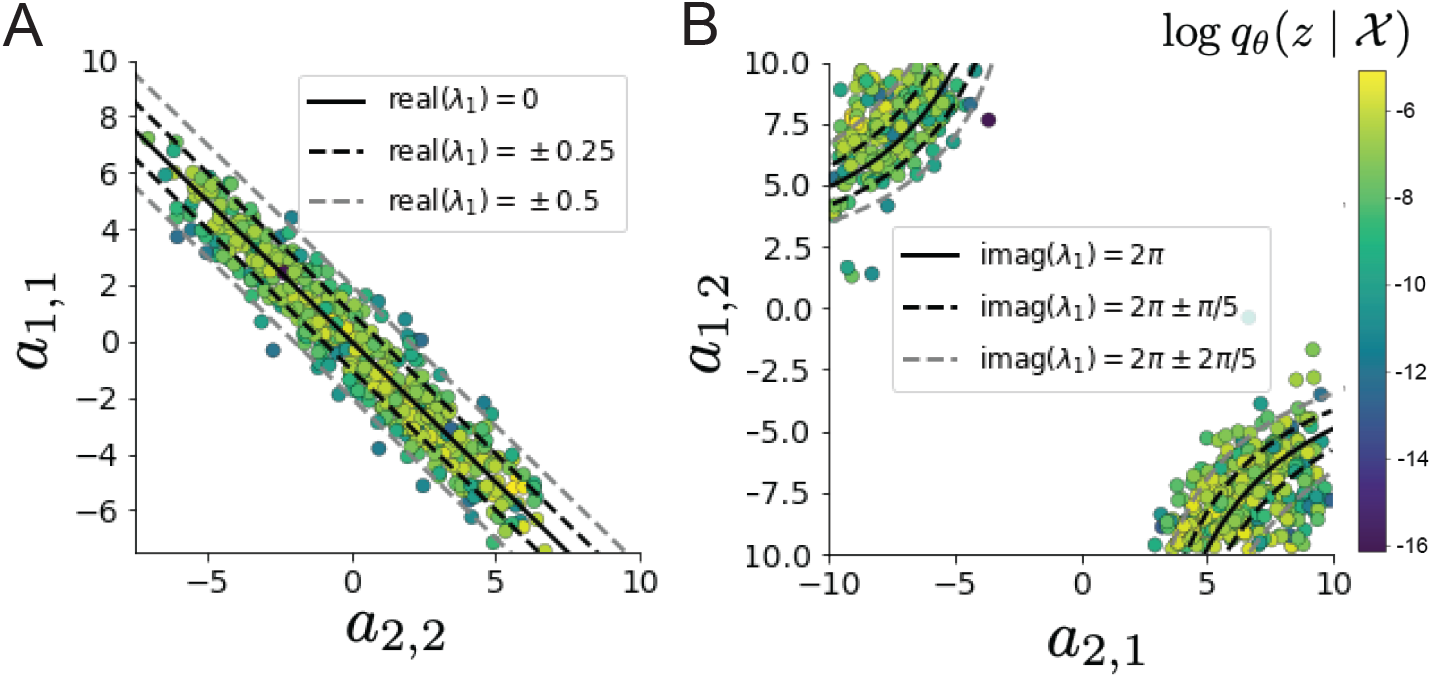
**A**. Probability contours in the *a*_1,1_-*a*_2,2_ plane were derived from the relationship to emergent property statistic of growth/decay factor real(*λ*_1_). **B**. Probability contours in the *a*_1,2_-*a*_2,1_ plane were derived from the emergent property statistic of oscillation frequency 2*π*imag(*λ*_1_).

**Figure S3:**
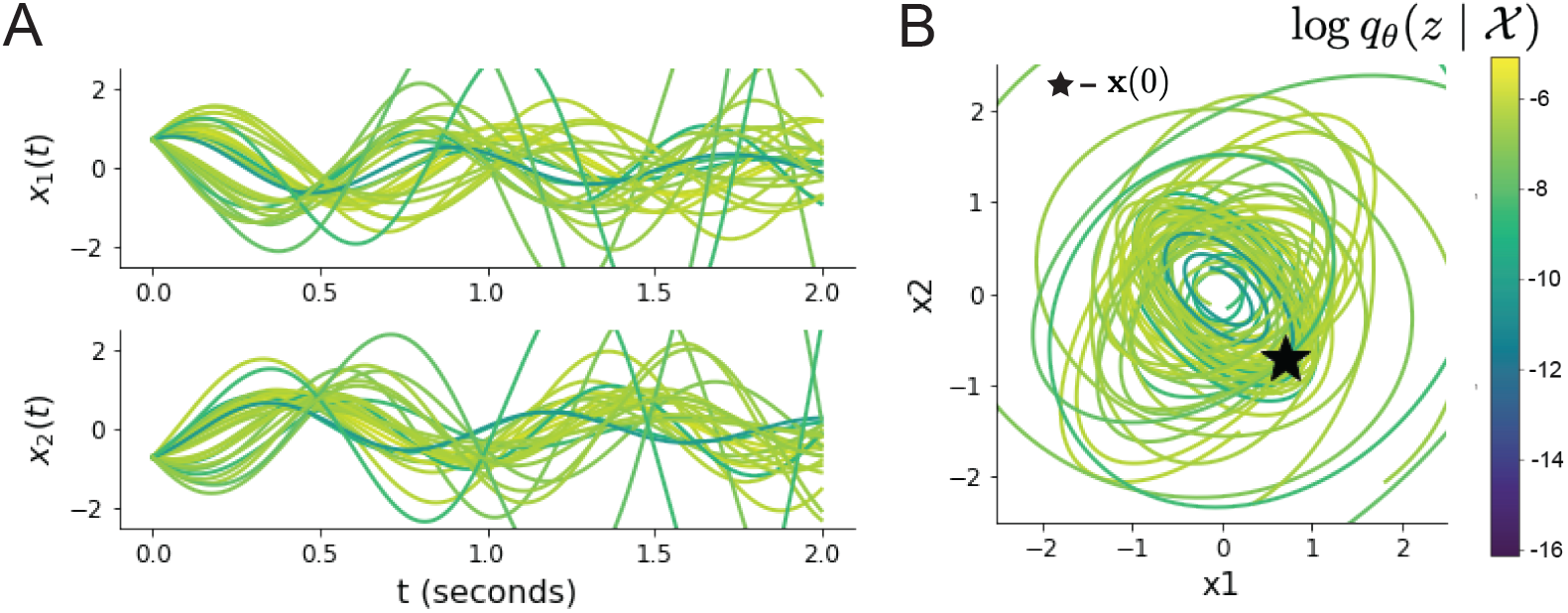
Sampled dynamical systems 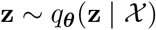 and their simulated activity from 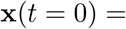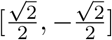 colored by log probability. **A**. Each dimension of the simulated trajectories throughout time. **B**. The simulated trajectories in phase space.

Since the marginal distribution of the data *p*(**x**) (or “evidence”) is independent of ***θ***, variational inference is executed by optimizing the remaining expression. This is usually framed as maximizing the evidence lower bound (ELBO)

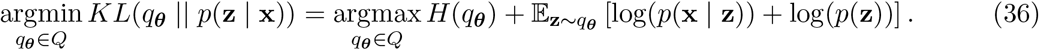

Now, we will show how the maximum entropy problem of EPI is equivalent to variational inference. In general, a maximum entropy problem (as in Equation 16) has an equivalent lagrange dual form:

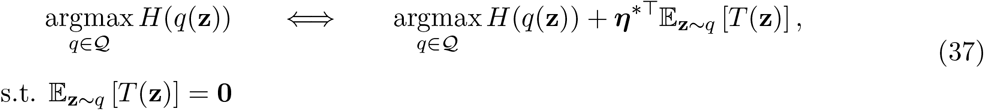

with lagrange multipliers ***η**\**. By moving the lagrange multipliers within the expectation

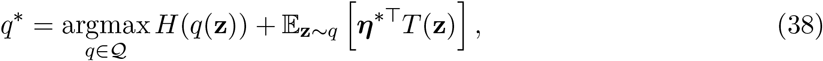

inserting a log exp(·) within the expectation,

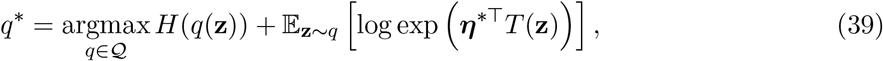

and finally choosing *T* (·) to be likelihood averaged statistics as in EPI

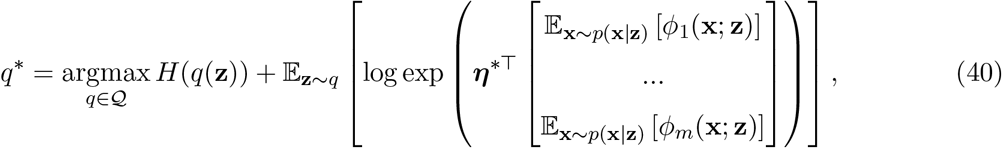

we can compare directly to the objective used in variational inference (Equation 36). We see that EPI is exactly variational inference with an exponential family likelihood defined by sufficient statistics 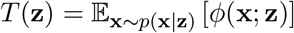, and where the natural parameter ***η**\** is predicated by the mean parameter ***μ***_opt_. Equation 40 implies that EPI uses an improper (or uniform) prior, which is easily changed.

This derivation of the equivalence between EPI and variational inference emphasizes why defining a statistical inference program by its mean parameterization ***μ***_opt_ is so useful. With EPI, one can clearly define the emergent property 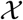 that the model of interest should produce through intuitive selection of ***μ***_opt_ for a given *T* (**z**). Alternatively, figuring out the correct natural parameters ***η**\** for the same *T*(**z**) that produces 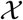 is a formally hard problem.

### 5.2 Stomatogastric ganglion

In Section 3.1 and 3.2, we used EPI to infer conductance parameters in a model of the stomatogastric ganglion (STG) [41]. This 5-neuron circuit model represents two subcircuits: that generating the pyloric rhythm (fast population) and that generating the gastric mill rhythm (slow population). The additional neuron (the IC neuron of the STG) receives inhibitory synaptic input from both subcircuits, and can couple to either rhythm dependent on modulatory conditions. There is also a parametric regime in which this neuron fires at an intermediate frequency between that of the fast and slow populations [41], which we infer with EPI as a motivational example. This model is not to be confused with an STG subcircuit model of the pyloric rhythm [68], which has been statistically inferred in other studies [15, 35].

#### 5.2.1 STG model

We analyze how the parameters **z** = [*g*_el_, *g*_synA_] govern the emergent phenomena of intermediate hub frequency in a model of the stomatogastric ganglion (STG) [41] shown in Figure 1A with activity **x** = [*x*_f1_, *x*_f2_, *x*_hub_, *x*_s1_, *x*_s2_], using the same hyperparameter choices as Gutierrez et al. Each neuron’s membrane potential *x_α_*(*t*) for *α* ∈ {f1, f2, hub, s1, s2} is the solution of the following stochastic differential equation:

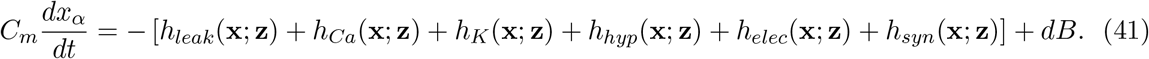

The input current of each neuron is the sum of the leak, calcium, potassium, hyperpolarization, electrical and synaptic currents. Each current component is a function of all membrane potentials and the conductance parameters **z**. Finally, we include gaussian noise *dB* to the model of Gutierrez et al. so that the model stochastic, although this is not required by EPI.

The capacitance of the cell membrane was set to *C_m_* = 1*nF*. Specifically, the currents are the difference in the neuron’s membrane potential and that current type’s reversal potential multiplied by a conductance:

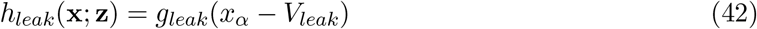

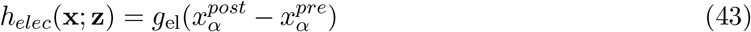

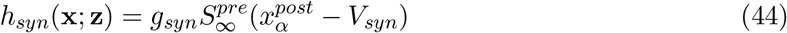

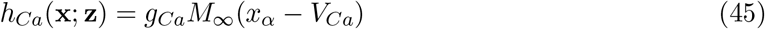

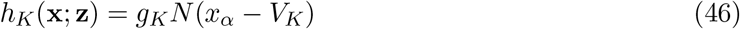

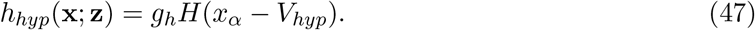

The reversal potentials were set to *V_leak_* = −40*mV*, *V_Ca_* = 100*mV*, *V_K_* = −80*mV*, *V_hyp_* = −20*mV*, and *V_syn_* = −75*mV*. The other conductance parameters were fixed to *g_leak_* = 1 × 10^*−*4^*μS*. *g_Ca_*, *g_K_*, and *g_hyp_* had different values based on fast, intermediate (hub) or slow neuron. The fast conductances had values *g_Ca_* = 1.9×10^*−*2^, *g_K_* = 3.9×10^*−*2^, and *g_hyp_* = 2.5×10^*−*2^. The intermediate conductances had values *g_Ca_* = 1.7 × 10^*−*2^, *g_K_* = 1.9 × 10^*−*2^, and *g_hyp_* = 8.0 × 10^*−*3^. Finally, the slow conductances had values *g_Ca_* = 8.5 × 10^*−*3^, *g_K_* = 1.5 × 10^*−*2^, and *g_hyp_* = 1.0 × 10^*−*2^.

Furthermore, the Calcium, Potassium, and hyperpolarization channels have time-dependent gating dynamics dependent on steady-state gating variables *M_∞_*, *N_∞_* and *H_∞_*, respectively:

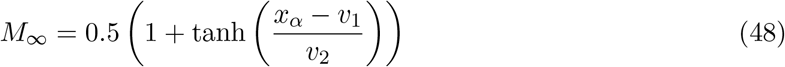

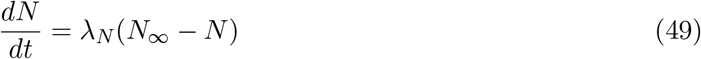

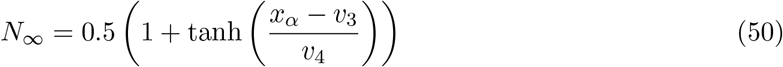

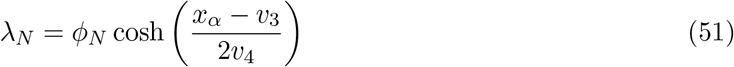

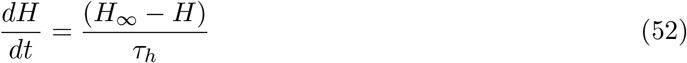

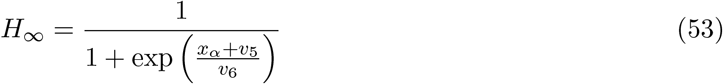

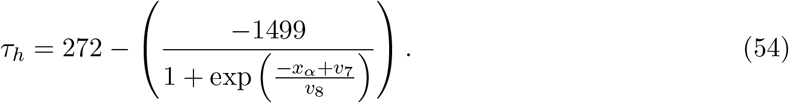

where we set *v*_1_ = 0*mV*, *v*_2_ = 20*mV*, *v*_3_ = 0*mV*, *v*_4_ = 15*mV*, *v*_5_ = 78.3*mV*, *v*_6_ = 10.5*mV*, *v*_7_ = −42.2*mV*, *v*_8_ = 87.3*mV*, *v*_9_ = 5*mV*, and *v_th_* = −25*mV*.

Finally, there is a synaptic gating variable as well:

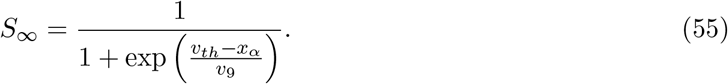

When the dynamic gating variables are considered, this is actually a 15-dimensional nonlinear dynamical system. The gaussian noise *d***B** has variance (1 × 10^*−*12^)^2^ A^2^, and introduces variability in frequency at each parameterization **z**.

#### 5.2.2 Hub frequency calculation

In order to measure the frequency of the hub neuron during EPI, the STG model was simulated for *T* = 300 time steps of *dt* = 25ms. The chosen *dt* and *T* were the most computationally convenient choices yielding accurate frequency measurement. We used a basis of complex exponentials with frequencies from 0.0-1.0 Hz at 0.01Hz resolution to measure frequency from simulated time series

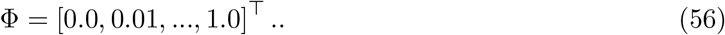

To measure spiking frequency, we processed simulated membrane potentials with a relu (spike extraction) and low-pass filter with averaging window of size 20, then took the frequency with the maximum absolute value of the complex exponential basis coefficients of the processed time-series. The first 20 temporal samples of the simulation are ignored to account for initial transients.

To differentiate through the maximum frequency identification, we used a soft-argmax Let 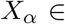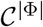 be the complex exponential filter bank dot products with the signal *x_α_* ∈ ℝ^*N*^, where *α* ∈ {f1, f2, hub, s1, s2}. The soft-argmax is then calculated using temperature parameter *β_ψ_* = 100

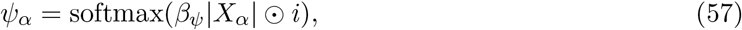

where *i* = [0, 1, *…,* 100]. The frequency is then calculated as

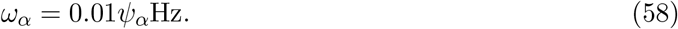

Intermediate hub frequency, like all other emergent properties in this work, is defined by the mean and variance of the emergent property statistics. In this case, we have one statistic, hub neuron frequency, where the mean was chosen to be 0.55Hz,(Equation 2) and variance was chosen to be 0.025^2^ Hz^2^ (Equation 3).

#### 5.2.3 EPI details for the STG model

EPI was run for the STG model using

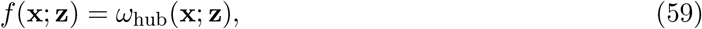

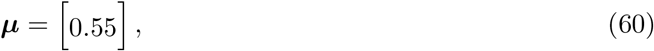

and

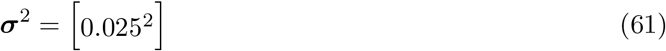

(see Sections 5.1.3–5.1.4, and example in Section 5.1.5). Throughout optimization, the augmented lagrangian parameters *η* and *c*, were updated after each epoch of *i*_max_ = 5, 000 iterations (see Section 5.1.4). The optimization converged after five epochs (Fig. S4).

**Figure S4:**
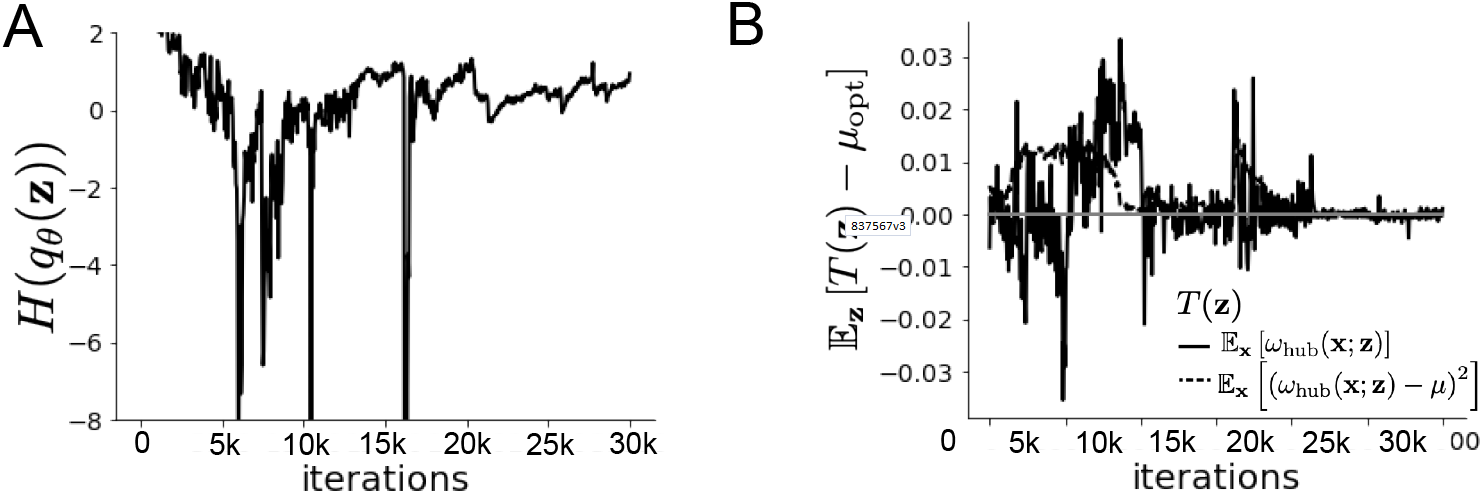
EPI optimization of the STG model producing network syncing. **A**. Entropy throughout optimization. **B**. The emergent property statistic means and variances converge to their constraints at 25,000 iterations following the fifth augmented lagrangian epoch.

For EPI in Fig 1E, we used a real NVP architecture with three coupling layers and two-layer neural networks of 25 units per layer. The normalizing flow architecture mapped 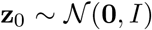 to a support of **z** = [*g*_el_, *g*_synA_] ∈ [4, 8] × [0.01, 4], initialized to a gaussian approximation of samples returned by a preliminary ABC search. We did not include *g*_synA_ < 0.01, for numerical stability. EPI optimization was run using 5 different random seeds for architecture initialization ***θ*** with an augmented lagrangian coefficient of *c*_0_ = 10^5^, *β* = 2, a batch size *n* = 400, and we simulated one **x**^(*i*)^ per **z**^(*i*)^. The architecture converged with criteria *N*_test_ = 100.

#### 5.2.4 Hessian sensitivity vectors

To quantify the second-order structure of the EPI distribution, we evaluated the Hessian of the log probability 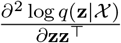. The eigenvector of this Hessian with most negative eigenvalue is defined as the sensitivity dimension **v**_1_, and all subsequent eigenvectors are ordered by increasing eigenvalue. These eigenvalues are quantifications of how fast the emergent property deteriorates via the parameter combination of their associated eigenvector. In Figure 1D, the sensitivity dimension *v*_1_ (solid) and the second eigenvector of the Hessian *v*_2_ (dashed) are shown evaluated at the mode of the distribution. Since the Hessian eigenvectors have sign degeneracy, the visualized directions in 2-D parameter space were chosen to have positive *g*_synA_. The length of the arrows is inversely proportional to the square root of the absolute value of their eigenvalues *λ*_1_ = −10.7 and *λ*_2_ = −3.22. For the same magnitude perturbation away from the mode, intermediate hub frequency only diminishes along the sensitivity dimension **v**_1_ (Fig. 1E-F).

### 5.3 Scaling EPI for stable amplification in RNNs

#### 5.3.1 Rank-2 RNN model

We examined the scaling properties of EPI by learning connectivities of RNNs of increasing size that exhibit stable amplification. Rank-2 RNN connectivity was modeled as *W* = *UV^⊤^*, where 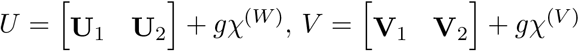, and 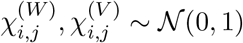. This RNN model has dynamics

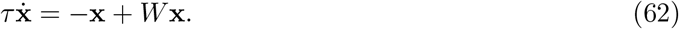

In this analysis, we inferred connectivity parameterizations 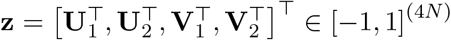 that produced stable amplification using EPI, SMC-ABC [26], and SNPE [35] (see Section Related Methods).

#### 5.3.2 Stable amplification

For this RNN model to be stable, all real eigenvalues of *W* must be less than 1: real(*λ*_1_) < 1, where *λ*_1_ denotes the greatest real eigenvalue of *W*. For a stable RNN to amplify at least one input pattern, the symmetric connectivity 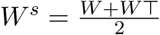 must have an eigenvalue greater than 1: 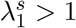, where *λ^s^* is the maximum eigenvalue of *W^s^*. These two conditions are necessary and sufficient for stable amplification in RNNs [51].

#### 5.3.3 EPI details for RNNs

We defined the emergent property of stable amplification with means of these eigenvalues (0.5 and 1.5, respectively) that satisfy these conditions. To complete the emergent property definition, we chose variances (0.25^2^) about those means such that samples rarely violate the eigenvalue constraints. To write the emergent property of Equation 5 in terms of the EPI optimization, we have

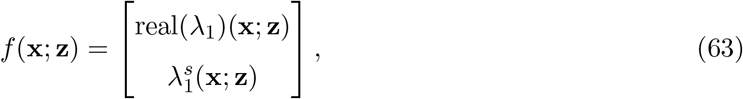

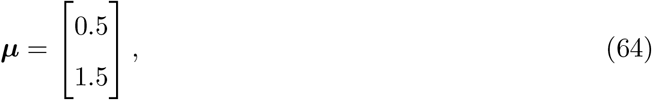

and

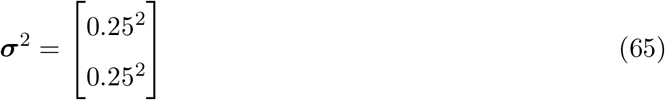

(see Sections 5.1.3–5.1.4, and example in Section 5.1.5). Gradients of maximum eigenvalues of Hermitian matrices like *W^s^* are available with modern automatic differentiation tools. To differentiate through the real(*λ*_1_), we solved the following equation for eigenvalues of rank-2 matrices using the rank reduced matrix *W^r^* = *V^⊤^U*

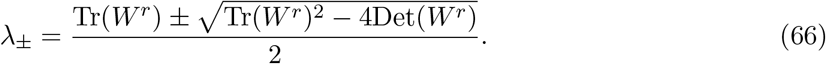

For EPI in Fig. 2, we used a real NVP architecture with three coupling layers of affine transformations parameterized by two-layer neural networks of 100 units per layer. The initial distribution was a standard isotropic gaussian 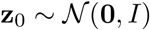 mapped to the support of **z**_*i*_ ∈ [−1, 1]. We used an augmented lagrangian coefficient of *c*_0_ = 10^3^, a batch size *n* = 200, *β* = 4, and we simulated one **W**^(*i*)^ per **z**^(*i*)^. We chose to use *i*_max_ = 500 iterations per augmented lagrangian epoch and emergent property constraint convergence was evaluated at *N*_test_ = 200 (Fig. 2B blue line, and Fig. 2C-D blue). It was fastest to initialize the EPI distribution on a Tesla V100 GPU, and then subsequently optimize it on a CPU with 32 cores. EPI timing measurements accounted for this initialization period.

#### 5.3.4 Methodological comparison

We compared EPI to two alternative simulation-based inference techniques, since the likelihood of these eigenvalues given **z** is not available. Approximate bayesian computation (ABC) [24] is a rejection sampling technique for obtaining sets of parameters **z** that produce activity **x** close to some observed data **x**_0_. Sequential Monte Carlo approximate bayesian computation (SMC-ABC) is the state-of-the-art ABC method, which leverages SMC techniques to improve sampling speed. We ran SMC-ABC with the pyABC package [94] to infer RNNs with stable amplification: connectivities having eigenvalues within an *ϵ*-defined *l*-2 distance of

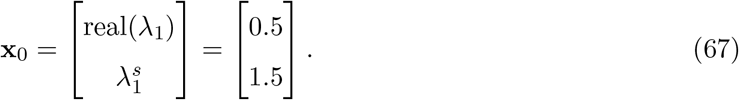

SMC-ABC was run with a uniform prior over **z** ∈ [−1, 1]^(4*N*)^, a population size of 1,000 particles with simulations parallelized over 32 cores, and a multivariate normal transition model.

**Figure S5:**
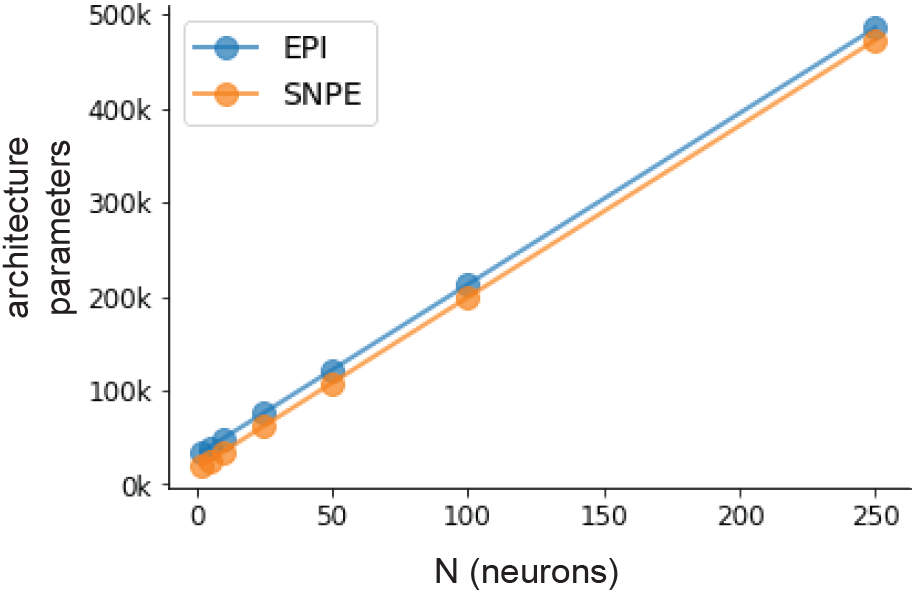
Number of parameters in deep probability distribution architectures of EPI (blue) and SNPE (orange) by RNN size (*N*).

SNPE, the next approach in our comparison, is far more similar to EPI. Like EPI, SNPE treats parameters in mechanistic models with deep probability distributions, yet the two learning algorithms are categorically different. SNPE uses a two-network architecture to approximate the posterior distribution of the model conditioned on observed data **x**_0_. The amortizing network maps observations **x**_*i*_ to the parameters of the deep probability distribution. The weights and biases of the parameter network are optimized by sequentially augmenting the training data with additional pairs (**z**_*i*_, **x**_*i*_) based on the most recent posterior approximation. This sequential procedure is important to get training data **z**_*i*_ to be closer to the true posterior, and **x**_*i*_ to be closer to the observed data. For the deep probability distribution architecture, we chose a masked autoregressive flow with affine couplings (the default choice), three transforms, 50 hidden units, and a normalizing flow mapping to the support as in EPI. This architectural choice closely tracked the size of the architecture used by EPI (Fig. S5). As in SMC-ABC, we ran SNPE with **x**_0_ = *μ*. All SNPE optimizations were run for a limit of 1.5 days, or until two consecutive rounds resulted in a validation log probability lower than the maximum observed for that random seed. It was always faster to run SNPE on a CPU with 32 cores rather than on a Tesla V100 GPU.

To compare the efficiency of these algorithms for inferring RNN connectivity distributions producing stable amplification, we develop a convergence criteria that can be used across methods. While EPI has its own hypothesis testing convergence criteria for the emergent property, it would not make sense to use this criteria on SNPE and SMC-ABC which do not constrain the means and variances of their predictions. Instead, we consider EPI and SNPE to have converged after completing its most recent optimization epoch (EPI) or round (SNPE) in which the distance 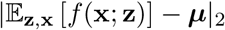 is less than 0.5. We consider SMC-ABC to have converged once the population produces samples within the *ϵ* = 0.5 ball ensuring stable amplification.

When assessing the scalability of SNPE, it is important to check that alternative hyperparamterizations could not yield better performance. Key hyperparameters of the SNPE optimization are the number of simulations per round *n*_round_, the number of atoms used in the atomic proposals of the SNPE-C algorithm [95], and the batch size *n*. To match EPI, we used a batch size of *n* = 200 for *N* <= 25, however we found *n* = 1, 000 to be helpful for SNPE in higher dimensions. While *n*_round_ = 1, 000 yielded SNPE convergence for *N* <= 25, we found that a substantial increase to *n*_round_ = 25, 000 yielded more consistent convergence at *N* = 50 (Fig. S6A). By increasing *n*_round_, we also necessarily increase the duration of each round. At *N* = 100, we tried two hyperparameter modifications. As suggested in [95], we increased *n*_atom_ by an order of magnitude to improve gradient quality, but this had little effect on the optimization (much overlap between same random seeds) (Fig. S6B). Finally, we increased *n*_round_ by an order of magnitude, which yielded convergence in one case, but no others. We found no way to improve the convergence rate of SNPE without making more aggressive hyperparameter choices requiring high numbers of simulations. In Figure 2C-D, we show samples from the random seed resulting in emergent property convergence at greatest entropy (EPI), the random seed resulting in greatest validation log probability (SNPE), and the result of all converged random seeds (SMC).

#### 5.3.5 Effect of RNN parameters on EPI and SNPE inferred distributions

To clarify the difference in objectives of EPI and SNPE, we show their results on RNN models with different numbers of neurons *N* and random strength *g*. The parameters inferred by EPI consistently produces the same mean and variance of real(*λ*_1_) and 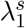, while those inferred by SNPE change according to the model definition (Fig. S7A). For *N* = 2 and *g* = 0.01, the SNPE posterior has greater concentration in eigenvalues around **x**_0_ than at *g* = 0.1, where the model has greater randomness (Fig. S7B top, orange). At both levels of *g* when *N* = 2, the posterior of SNPE has lower entropy than EPI at convergence (Fig. S7B top). However at *N* = 10, SNPE results in a predictive distribution of more widely dispersed eigenvalues (Fig. S7A bottom), and an inferred posterior with greater entropy than EPI (Fig. S7B bottom). We highlight these differences not to focus on an insightful trend, but to emphasize that these methods optimize different objectives with different implications.

**Figure S6:**
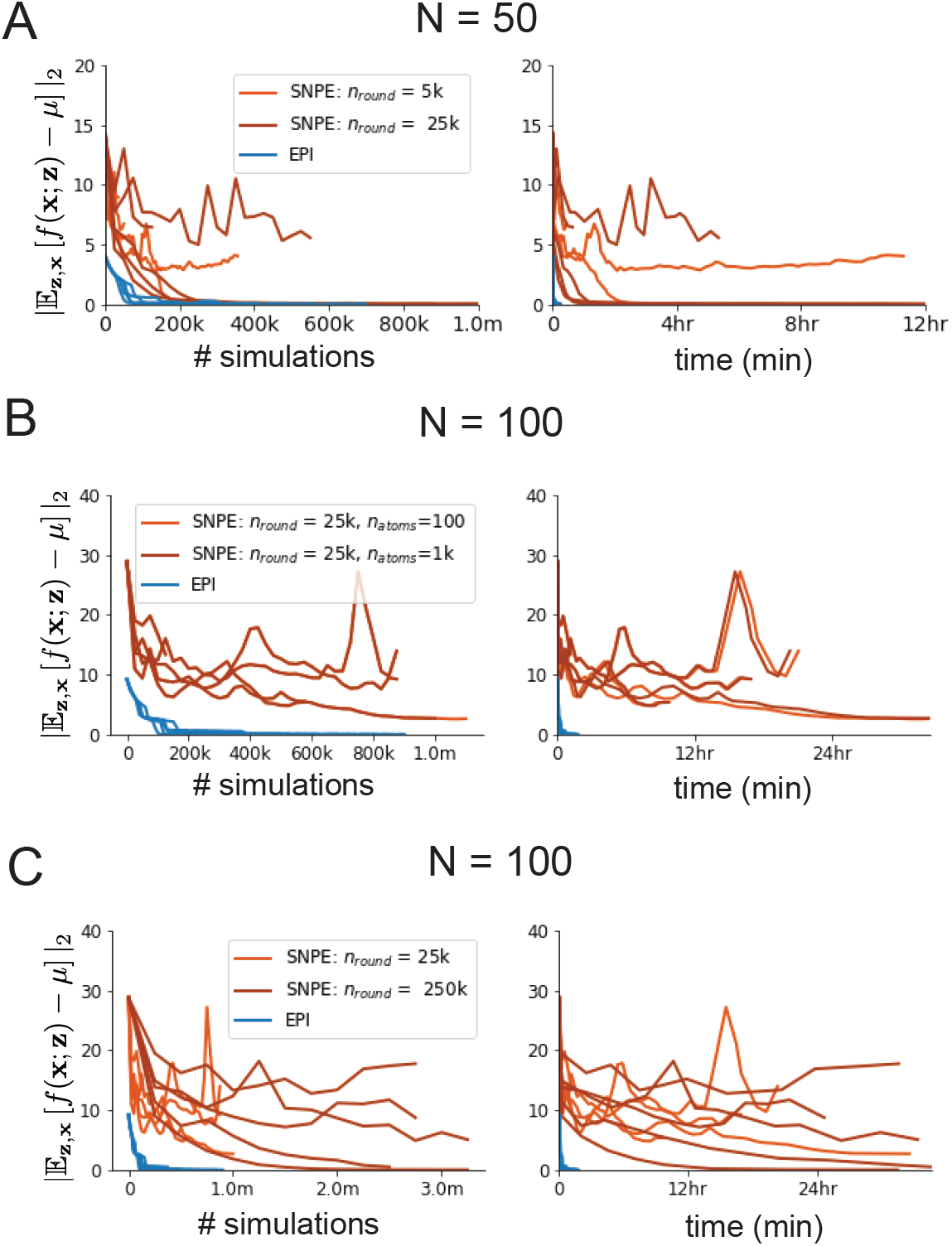
SNPE convergence was enabled by increasing *n*_round_, not *n*_atom_. **A**. Difference of mean predictions **x**_0_ throughout optimization at *N* = 50 with by simulation count (left) and wall time (right) of SNPE with *n*_round_ = 5, 000 (light orange), SNPE with *n*_round_ = 25, 000 (dark orange), and EPI (blue). Each line shows an individual random seed. **B**. Same conventions as A at *N* = 100 of SNPE with *n*_atom_ = 100 (light orange) and *n*_atom_ = 1, 000 (dark orange). **C**. Same conventions as A at *N* = 100 of SNPE with *n*_round_ = 25, 000 (light orange) and *n*_round_ = 250, 000 (dark orange).

Note that SNPE converges when it’s validation log probability has saturated after several rounds of optimization (Fig. S7C), and that EPI converges after several epochs of its own optimization to enforce the emergent property constraints (Fig. S7D blue). Importantly, as SNPE optimizes its posterior approximation, the predictive means change, and at convergence may be different than **x**_0_ (Fig. S7D orange, left). It is sensible to assume that predictions of a well-approximated SNPE posterior should closely reflect the data on average (especially given a uniform prior and a low degree of stochasticity), however this is not a given. Furthermore, no aspect of the SNPE optimization controls the variance of the predictions (Fig. S7D orange, right).

### 5.4 Primary visual cortex

#### 5.4.1 V1 model

E-I circuit models, rely on the assumption that inhibition can be studied as an indivisible unit, despite ample experimental evidence showing that inhibition is instead composed of distinct elements [63]. In particular three types of genetically identified inhibitory cell-types – parvalbumin (P), somatostatin (S), VIP (V) – compose 80% of GABAergic interneurons in V1 [61–63], and follow specific connectivity patterns (Fig. 3A) [64], which lead to cell-type specific computations [47, 96]. Currently, how the subdivision of inhibitory cell-types, shapes correlated variability by reconfiguring recurrent network dynamics is not understood.

In the stochastic stabilized supralinear network [59], population rate responses **x** to mean input **h**, recurrent input *W***x** and slow noise ***ϵ*** are governed by

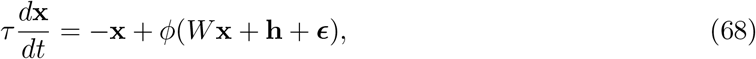

where the noise is an Ornstein-Uhlenbeck process ***ϵ*** ~ *OU* (*τ*_noise_, ***σ***)

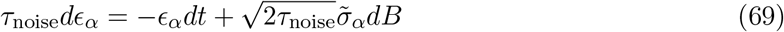

with *τ*_noise_ = 5ms > *τ* = 1ms. The noisy process is parameterized as

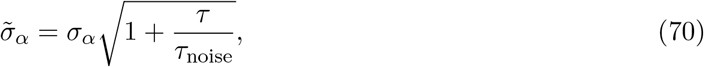

so that ***σ*** parameterizes the variance of the noisy input in the absence of recurrent connectivity (*W* = **0**). As contrast *c* ∈ [0, 1] increases, input to the E- and P-populations increases relative to a baseline input **h** = **h**_*b*_ + *c***h**_*c*_. Connectivity (*W*_fit_) and input (**h**_*b,*fit_ and **h**_*c,*fit_) parameters were fit using the deterministic V1 circuit model [47]

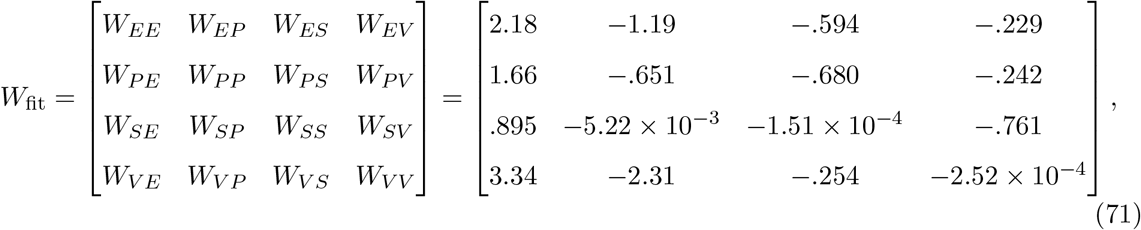

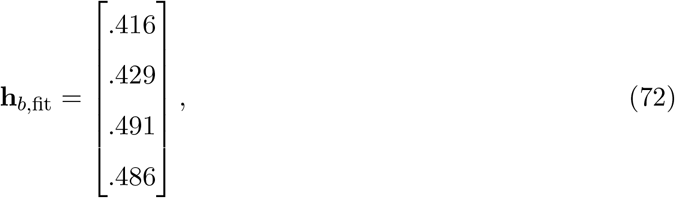

and

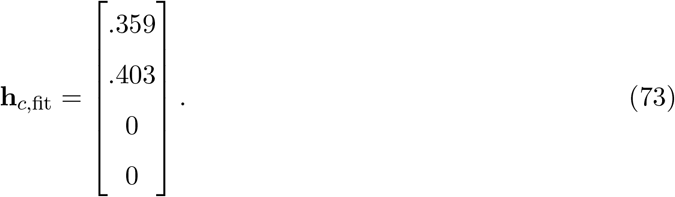

**Figure S7:**
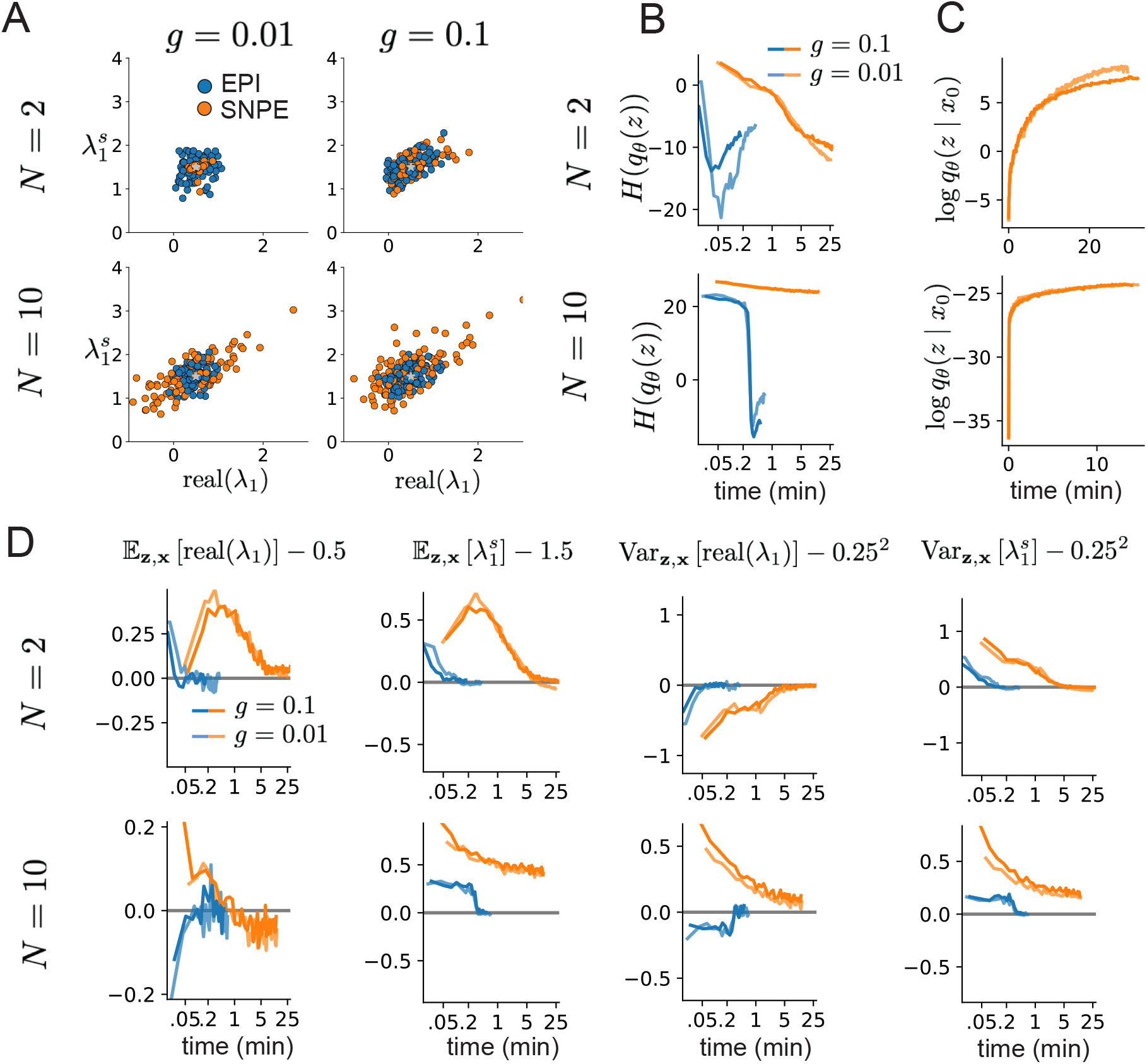
Model characteristics affect predictions of posteriors inferred by SNPE, while predictions of parameters inferred by EPI remain fixed. **A**. Predictive distribution of EPI (blue) and SNPE (orange) inferred connectivity of RNNs exhibiting stable amplification with *N* = 2 (top), *N* = 10 (bottom), *g* = 0.01 (left), and *g* = 0.1 (right). **B**. Entropy of parameter distribution approximations throughout optimization with *N* = 2 (top), *N* = 10 (bottom), *g* = 0.1 (dark shade), and *g* = 0.01 (light shade). **C**. Validation log probabilities throughout SNPE optimization. Same conventions as B. **D**. Adherence to EPI constraints. Same conventions as B.

To obtain rates on a realistic scale (100-fold greater), we map these fitted parameters to an equivalence class

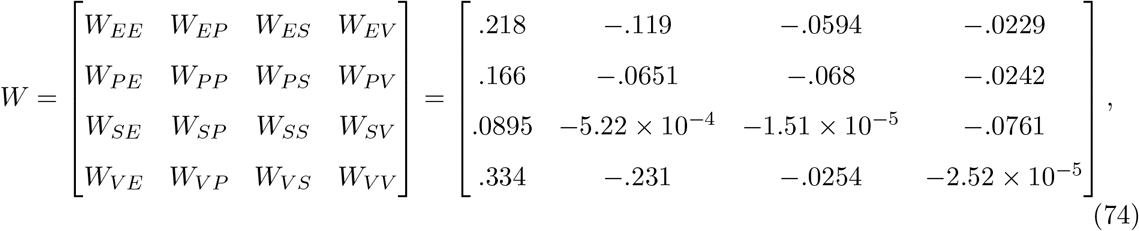

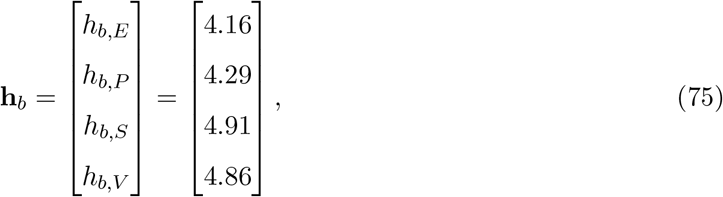

and

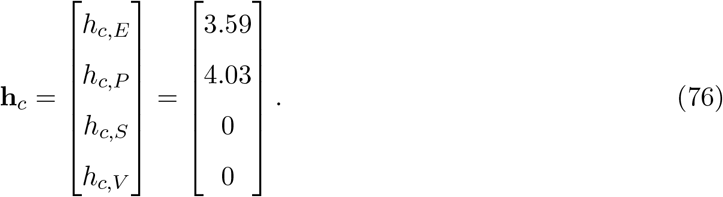

Circuit responses are simulated using *T* = 200 time steps at *dt* = 0.5ms from an initial condition drawn from **x**(0) ~ *U* [10Hz, 25Hz]. Standard deviation of the E-population *s_E_*(**x**; **z**) is calculated as the square root of the temporal variance from *t_ss_* = 75ms to *Tdt* = 100ms

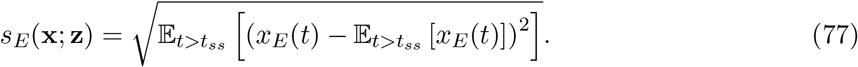

#### 5.4.2 EPI details for the V1 model

To write the emergent properties of Equation 7 in terms of the EPI optimization, we have

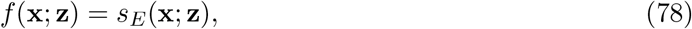

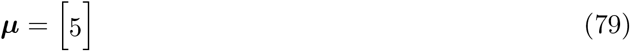

(or ***μ*** = [10]), and

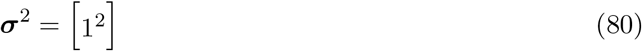

(see Sections 5.1.3–5.1.4, and example in Section 5.1.5).

For EPI in Figures 3D-E and S8, we used a real NVP architecture with three coupling layers and two-layer neural networks of 50 units per layer. The normalizing flow architecture mapped 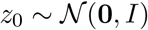 to a support of **z** = [*σ_E_, σ_P_, σ_S_, σ_V_*] ∈ [0.0, 0.5]^4^. EPI optimization was run using three different random seeds for architecture initialization ***θ*** with an augmented lagrangian coefficient of *c*_0_ = 10^*−*1^, *β* = 2, a batch size *n* = 100, and simulated 100 trials to calculate average *s_E_*(**x**; **z**) for each **z**^(*i*)^. We used *i*_max_ = 2, 000 iterations per epoch. The distributions shown are those of the architectures converging with criteria *N*_test_ = 100 at greatest entropy across three random seeds. Optimization details are shown in Figure S9. The sums of squares of each pair of parameters are shown for each EPI distribution in Figure S10. The plots are histograms of 500 samples from each EPI distribution from which the significance *p*-values of Section 3.4 are determined.

**Figure S8:**
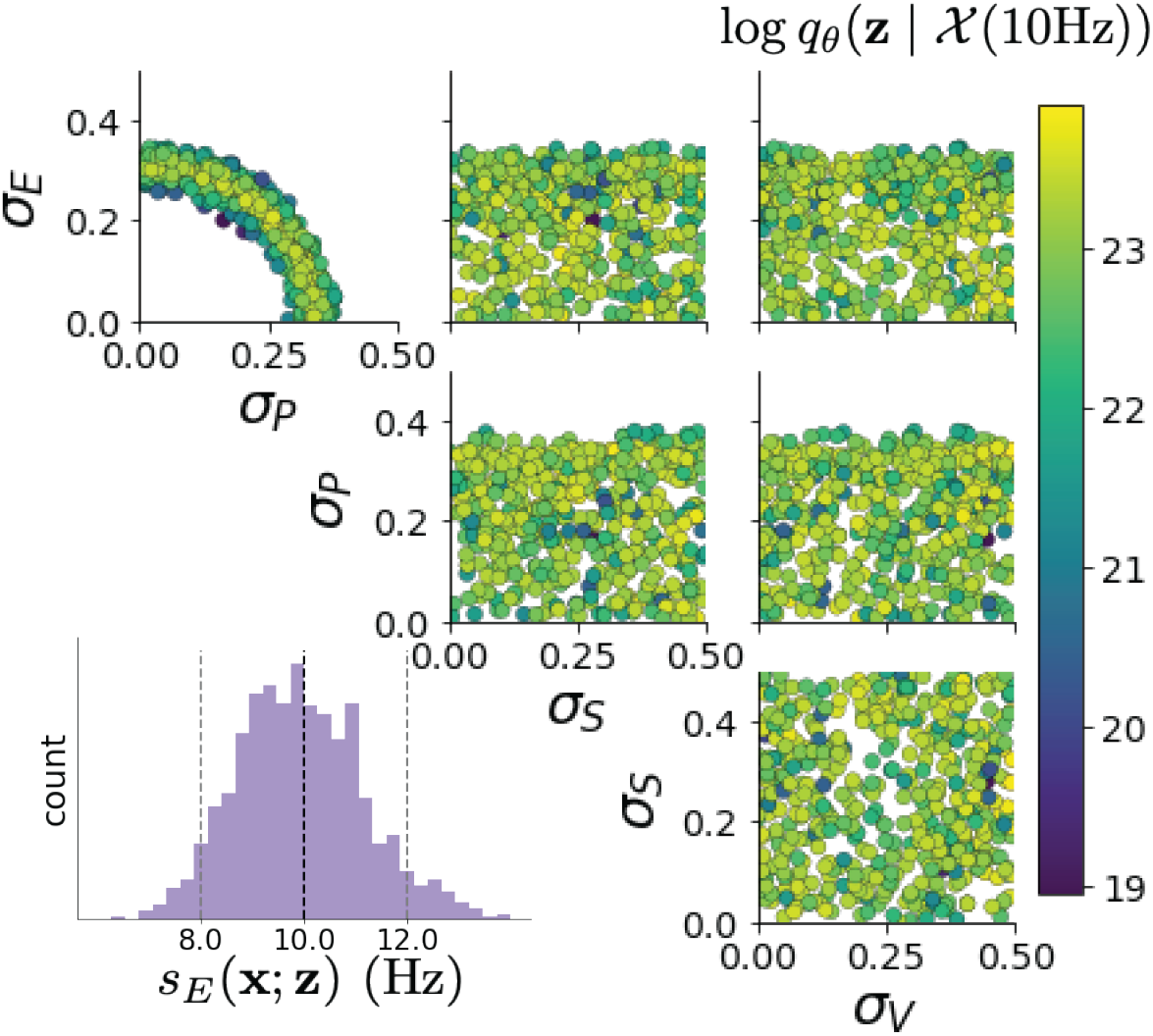
EPI inferred distribution for 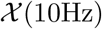.

#### 5.4.3 Sensitivity analyses

In Fig. 3E, we visualize the modes of 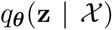 throughout the *σ_E_*-*σ_P_* marginal. At each local mode **z**\* (*σ_P_*), where *σ_P_* is fixed, we calculated the Hessian and visualized the sensitivity dimension in the direction of positive *σ_E_*.

#### 5.4.4 Testing for the paradoxical effect

The paradoxical effect occurs when a populations steady state rate is decreased (or increased) when an increase (decrease) in current is applied to that population [12]. To see which, if any, populations exhibited a paradoxical effect, we examined responses to changes in input (Fig. S11). Input magnitudes were chosen so that the effect is salient (0.002 for E and P, but 0.02 for S and V). Only the P-population exhibited the paradoxical effect at this connectivity *W* and input **h**.

#### 5.4.5 Primary visual cortex: Mathematical intuition and challenges

The dynamical system that we are working with can be written as

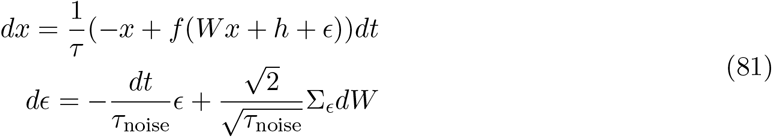

**Figure S9:**
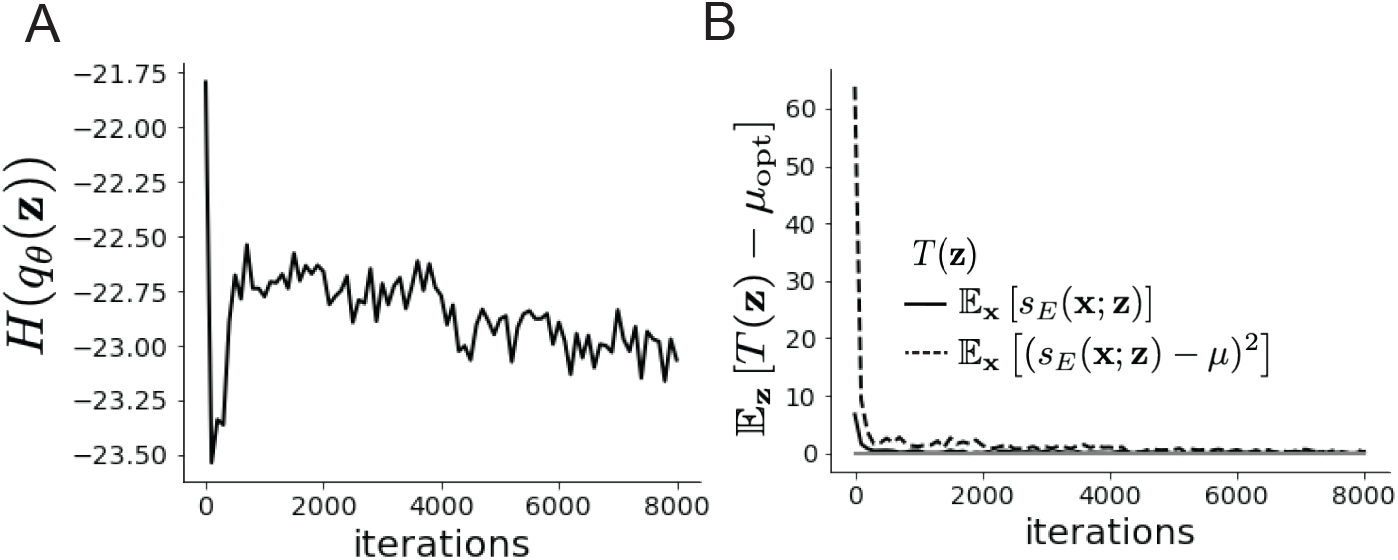
EPI optimization 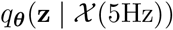 **A**. Entropy throughout optimization. **B**. The emergent property statistic means and variances converge to their constraints at 8,000 iterations following the fourth augmented lagrangian epoch.

Where in this paper we chose

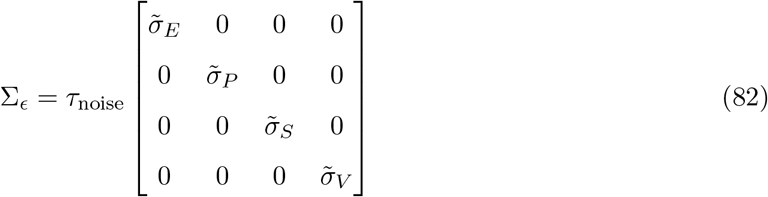

where 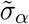 is the reparameterized standard deviation of the noise for population *α* from Equation 70.

In order to compute this covariance, we define *v* = *ωx* + *h* + *ϵ* and *S* = *I* − *ωf′*(*v*)), to re-write Eq. (81) as an 8-dimensional system:

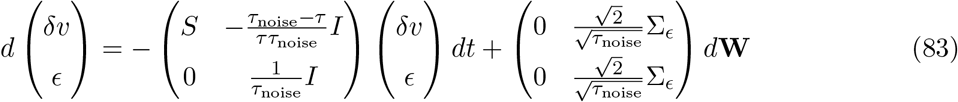

Where *d***W** is a vector with the private noise of each variable. The *d***W** term is multiplied by a non-diagonal matrix is because the noise that the voltage receives is the exact same than the one that comes from the OU process and not another process. The solution of this problem is given by the Lyapunov Equation [59, 66]:

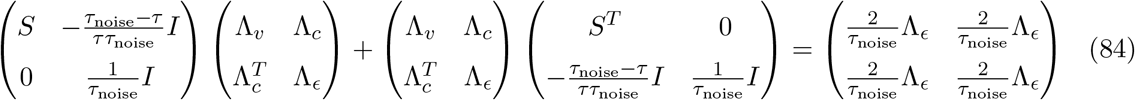

**Figure S10:**
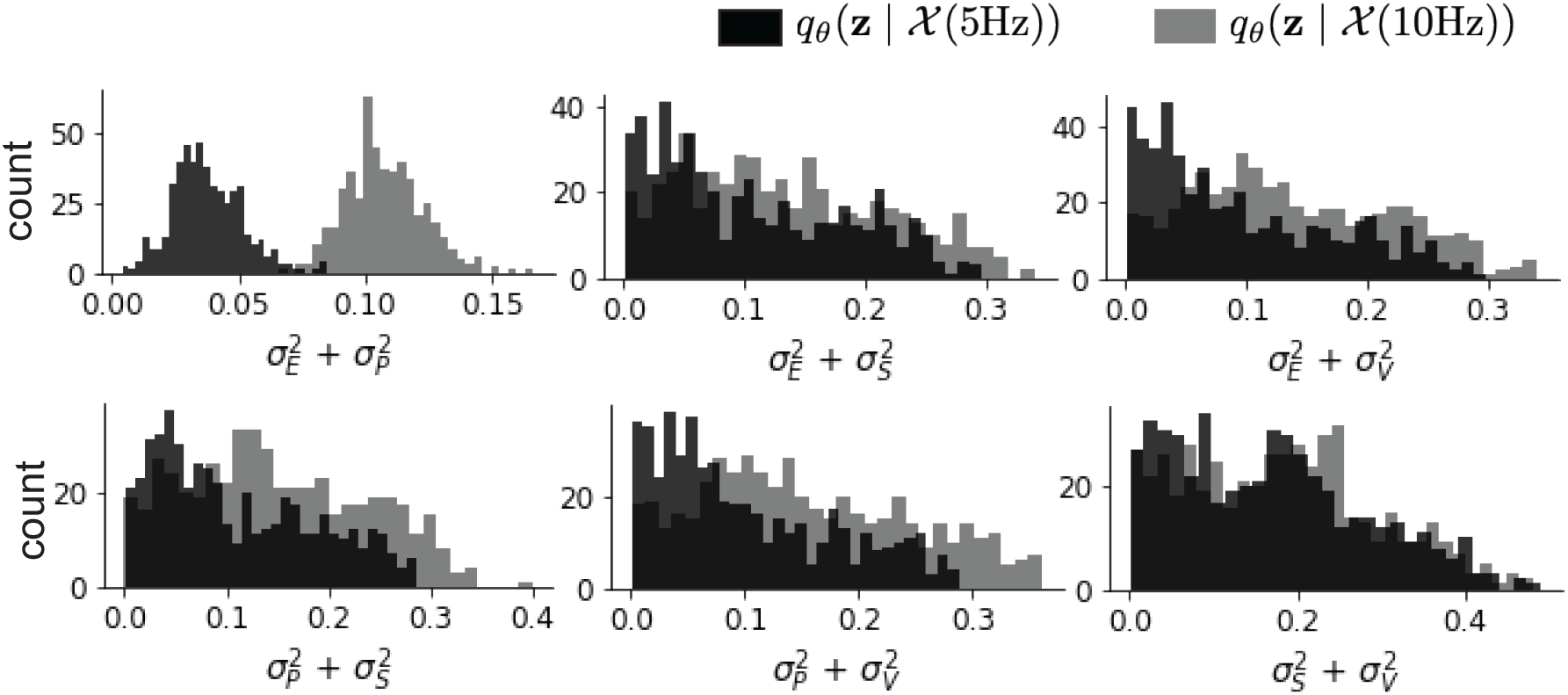
EPI predictive distributions of the sum of squares of each pair of noise parameters.

To obtain an equation for Λ_*v*_, we solve this block matrix multiplication:

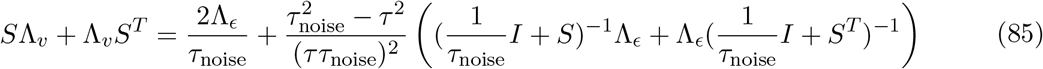

Which is another Lyapunov Equation, now in 4 dimensions. In the simplest case in which *τ*_noise_ = *τ*, the voltage is directly driven by white noise, and Λ_*v*_ can be expressed in powers of *S* and *S^T^*. Because *S* satisfies its own polynomial equation (Cayley Hamilton theorem), there will be 4 coefficients for the expansion of *S* and 4 for *S^T^*, resulting in 16 coefficients that define Λ_*v*_ for a given *S*. Due to symmetry arguments [66], in this case the diagonal elements of the covariance matrix of the voltage will have the form:

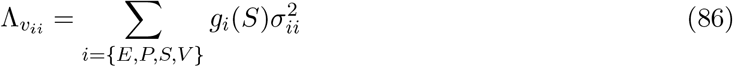

These coefficients *g_i_*(*S*) are complicated functions of the Jacobian of the system. Although expressions for these coefficients can be found explicitly, only numerical evaluation of those expressions determine which components of the noisy input are going to strongly influence the variability of excitatory population. Showing the generality of this dependence in more complicated noise scenarios (e.g. *τ*_noise_ > *τ* as in Section 3.4), is the focus of current research.

**Figure S11:**
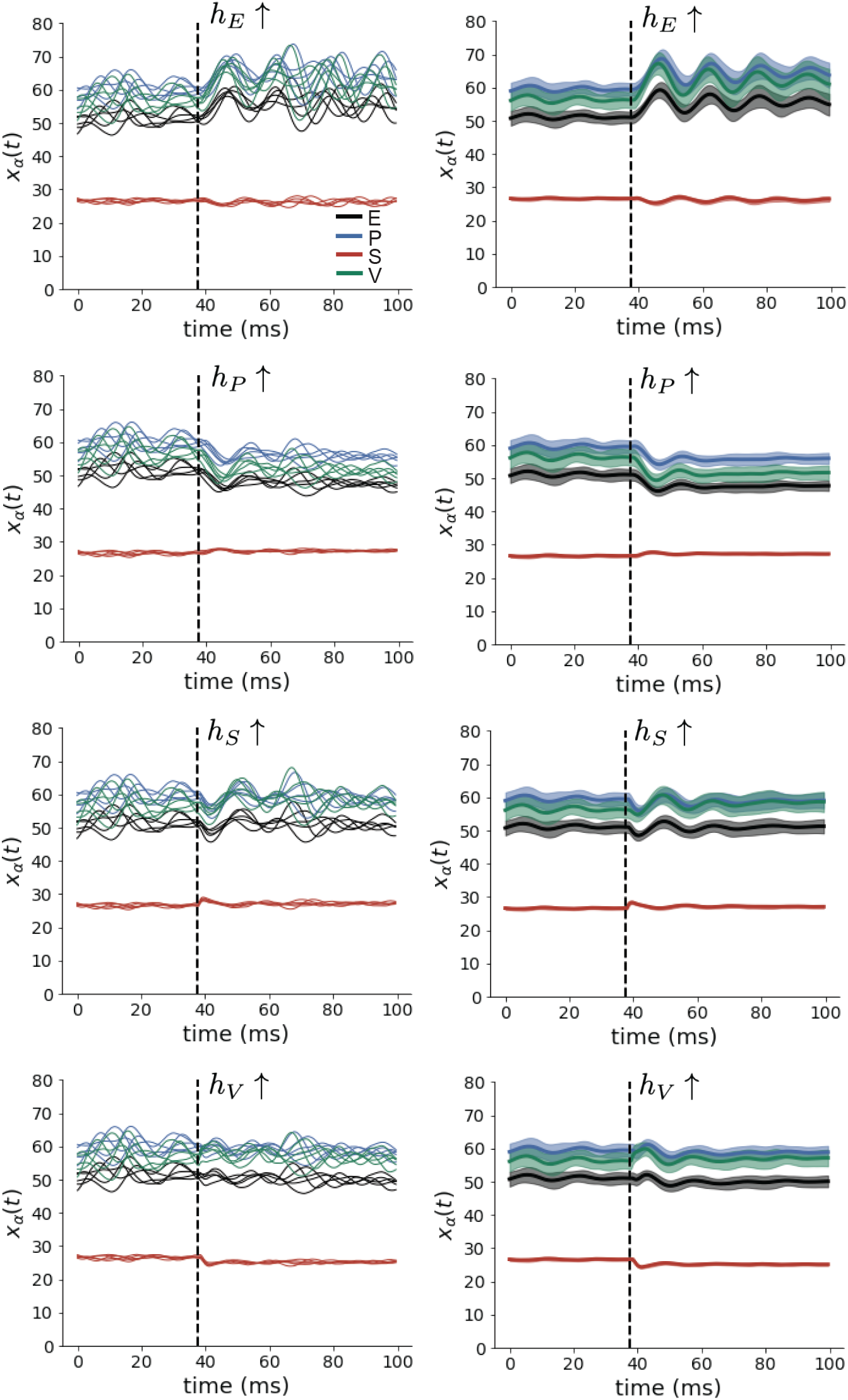
(Left) SSSN simulations for small increases in neuron-type population input. (Right) Average (solid) and standard deviation (shaded) of stochastic fluctuations of responses.

### 5.5 Superior colliculus

#### 5.5.1 SC model

The ability to switch between two separate tasks throughout randomly interleaved trials, or “rapid task switching,” has been studied in rats, and midbrain superior colliculus (SC) has been show to play an important in this computation [67]. Neural recordings in SC exhibited two populations of neurons that simultaneously represented both task context (Pro or Anti) and motor response (contralateral or ipsilateral to the recorded side), which led to the distinction of two functional classes: the Pro/Contra and Anti/Ipsi neurons [48]. Given this evidence, Duan et al. proposed a model with four functionally-defined neuron-type populations: two in each hemisphere corresponding to the Pro/Contra and Anti/Ipsi populations. We study how the connectivity of this neural circuit governs rapid task switching ability.

The four populations of this model are denoted as left Pro (LP), left Anti (LA), right Pro (RP) and right Anti (RA). Each unit has an activity (*x_α_*) and internal variable (*u_α_*) related by

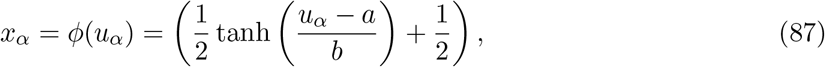

where *α* ∈ {*LP, LA, RA, RP*}, *a* = 0.05 and *b* = 0.5 control the position and shape of the nonlinearity. We order the neural populations of *x* and *u* in the following manner

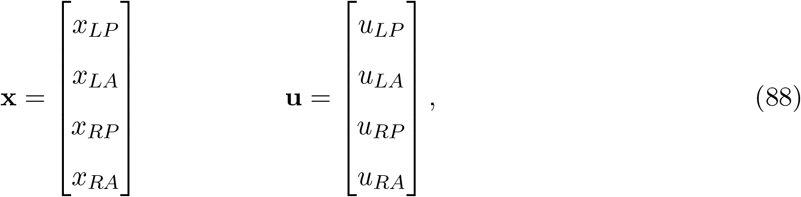

which evolve according to

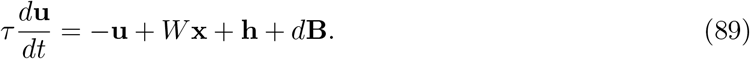

with time constant *τ* = 0.09*s*, step size 24ms and Gaussian noise *d***B** of variance 0.2^2^. These hyperparameter values are motivated by modeling choices and results from [48].

The weight matrix has 4 parameters for self *sW*, vertical *vW*, horizontal *hW*, and diagonal *dW* connections:

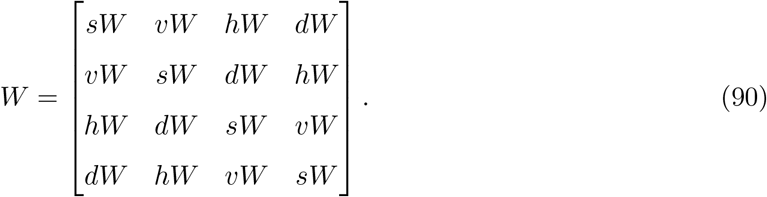

**Figure S12:**
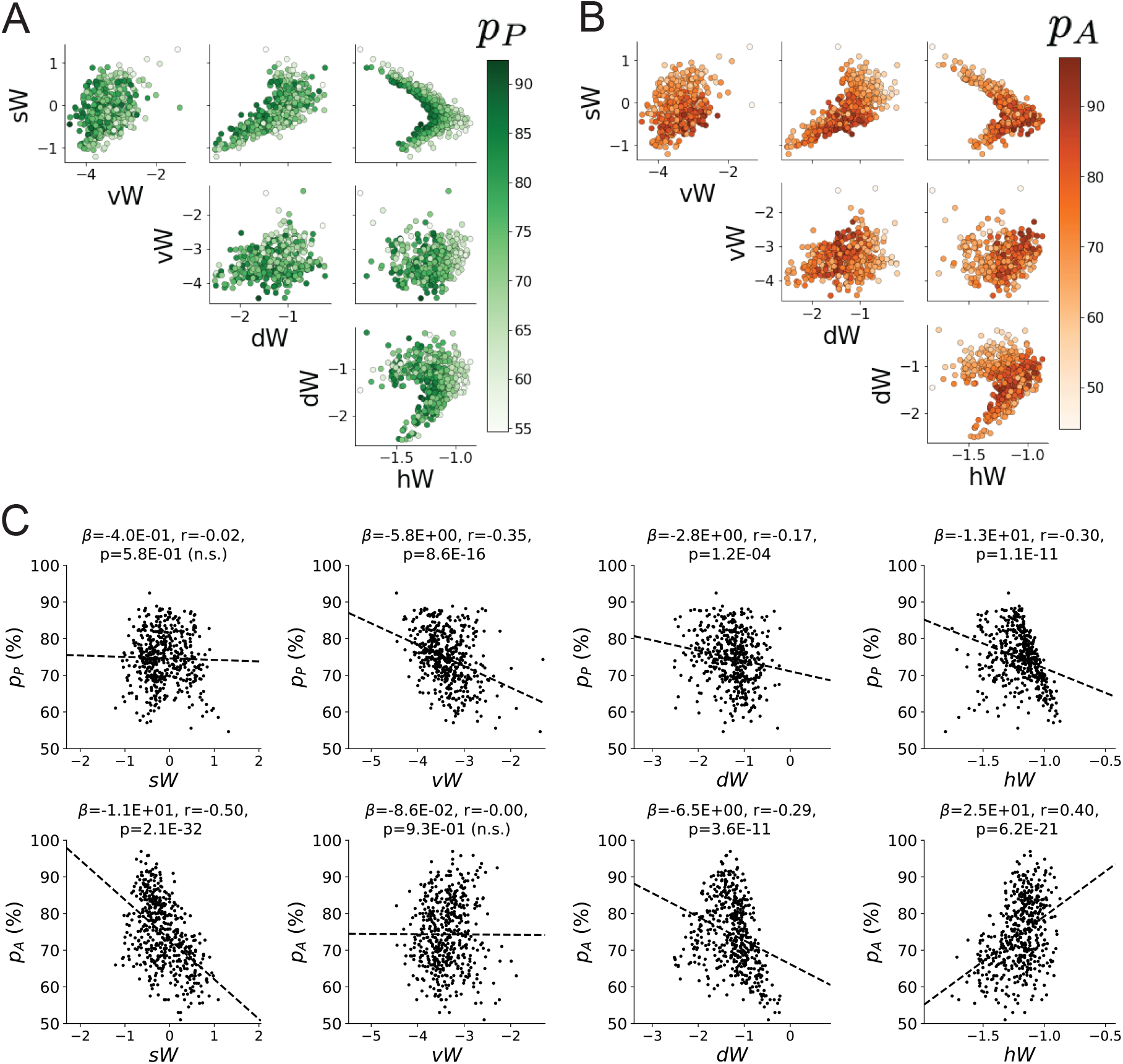
**A**. Same pairplot as Fig. 4C colored by Pro task accuracy. **B**. Same as A colored by Anti task accuracy. **C**. Connectivity parameters of EPI distributions versus task accuracies. *β* is slope coefficient of linear regression, *r* is correlation, and *p* is the two-tailed p-value.

**Figure S13:**
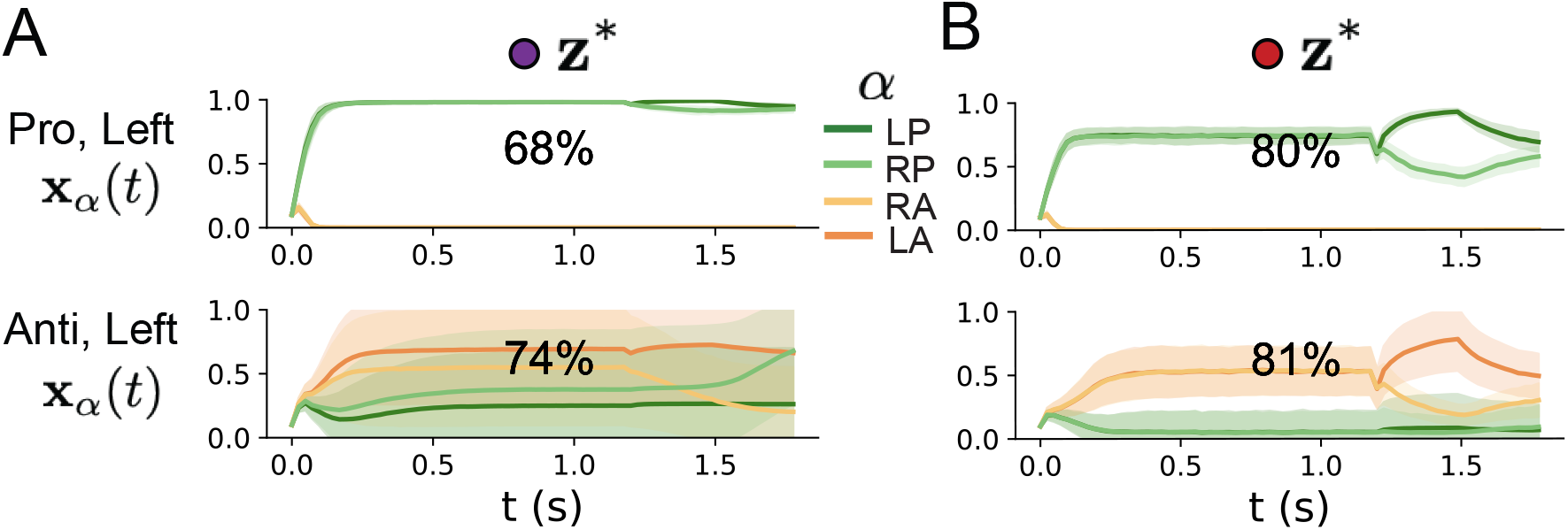
**A**. Simulations in network regime 1: **z**\* (*sW* = −0.75). **B**. Simulations in network regime 2: **z**\* (*sW* = 0.75).

We study the role of parameters **z** = [*sW, vW, hW, dW*]^⊤^ in rapid task switching.

The circuit receives four different inputs throughout each trial, which has a total length of 1.8s.

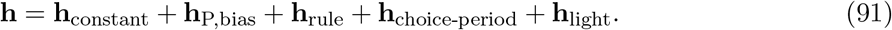

There is a constant input to every population,

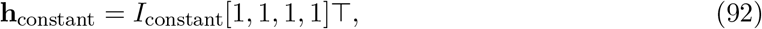

a bias to the Pro populations

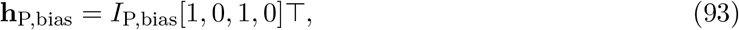

rule-based input depending on the condition

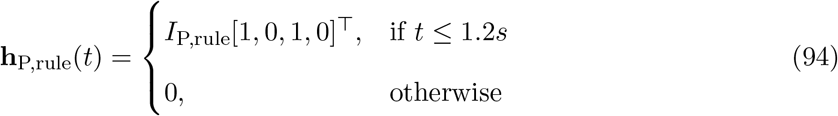

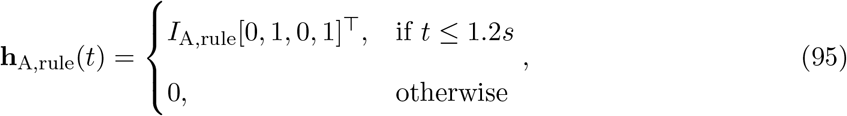

a choice-period input

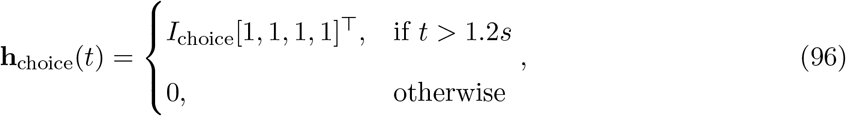

and an input to the right or left-side depending on where the light stimulus is delivered

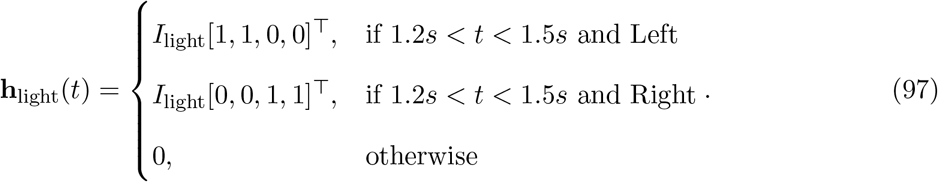

**Figure S14:**
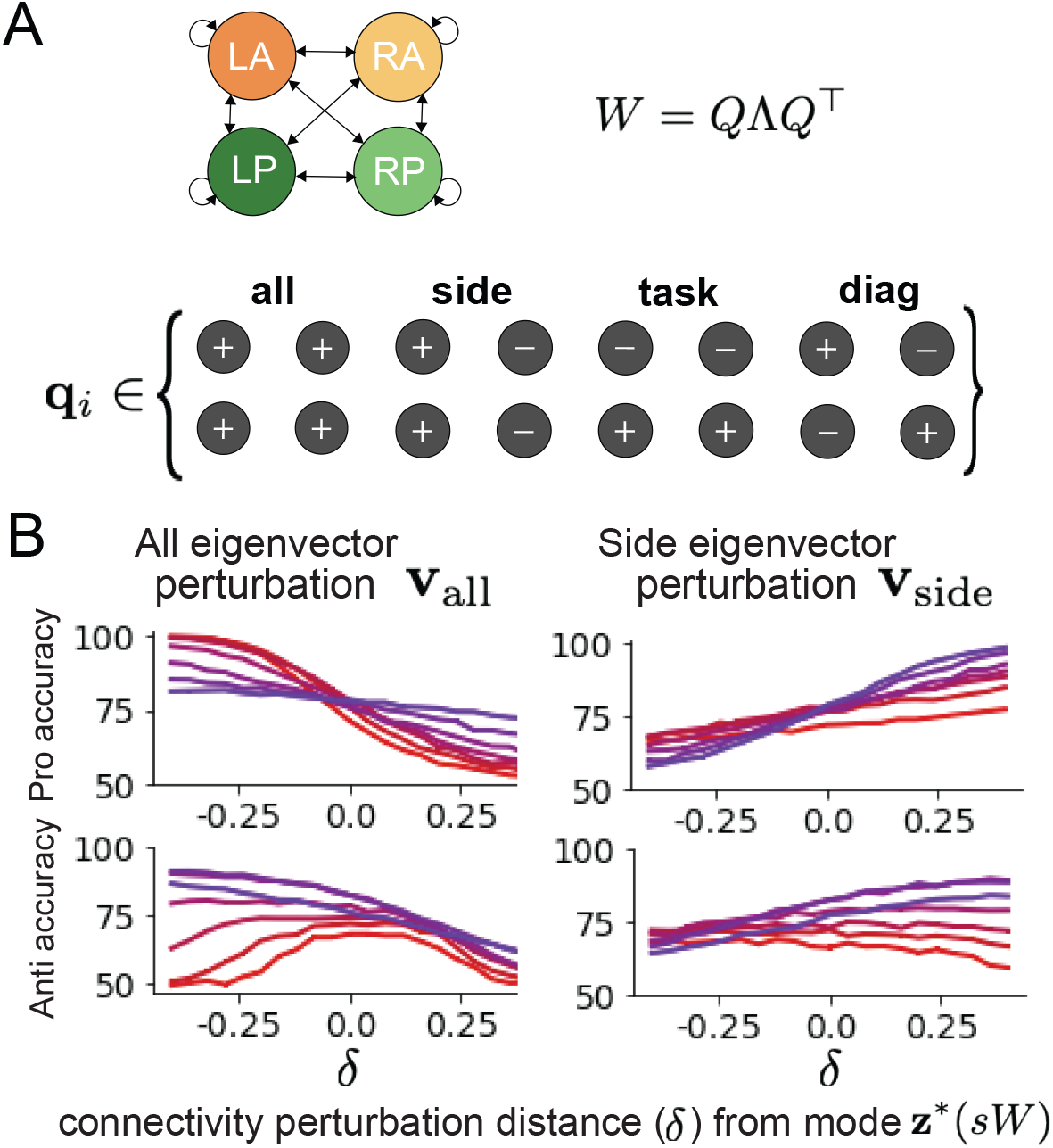
**A**. Invariant eigenvectors of connectivity matrix *W*. **B**. Accuracies for connectivity perturbations when changing *λ*_all_ and *λ*_side_ (*λ*_task_ and *λ*_diag_ shown in Fig. 4D).

The input parameterization was fixed to *I*_constant_ = 0.75, *I*_P,bias_ = 0.5, *I*_P,rule_ = 0.6, *I*_A,rule_ = 0.6, *I*_choice_ = 0.25, and *I*_light_ = 0.5.

#### 5.5.2 Task accuracy calculation

The accuracies of the Pro and Anti tasks are calculated as

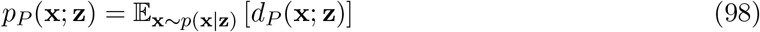

and

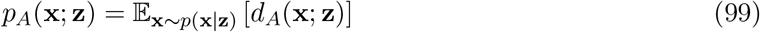

where *d_P_* (**x**; **z**) and *d_A_*(**x**; **z**) calculate the decision made in each trial (approximately 1 for correct and 0 for incorrect choices). Specifically,

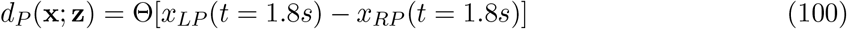

in Pro trials where the stimulus is on the left side, and Θ approximates the Heaviside step function. Similarly,

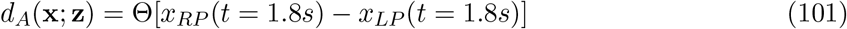

in Anti trials where the stimulus was on the left side. Our accuracy calculation only considers one stimulus presentation (Left), since the model is left-right symmetric. The accuracy is averaged over 200 independent trials, and the Heaviside step function is approximated as

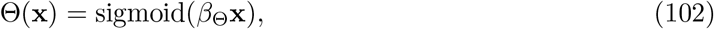

where *β*_Θ_ = 100.

**Figure S15:**
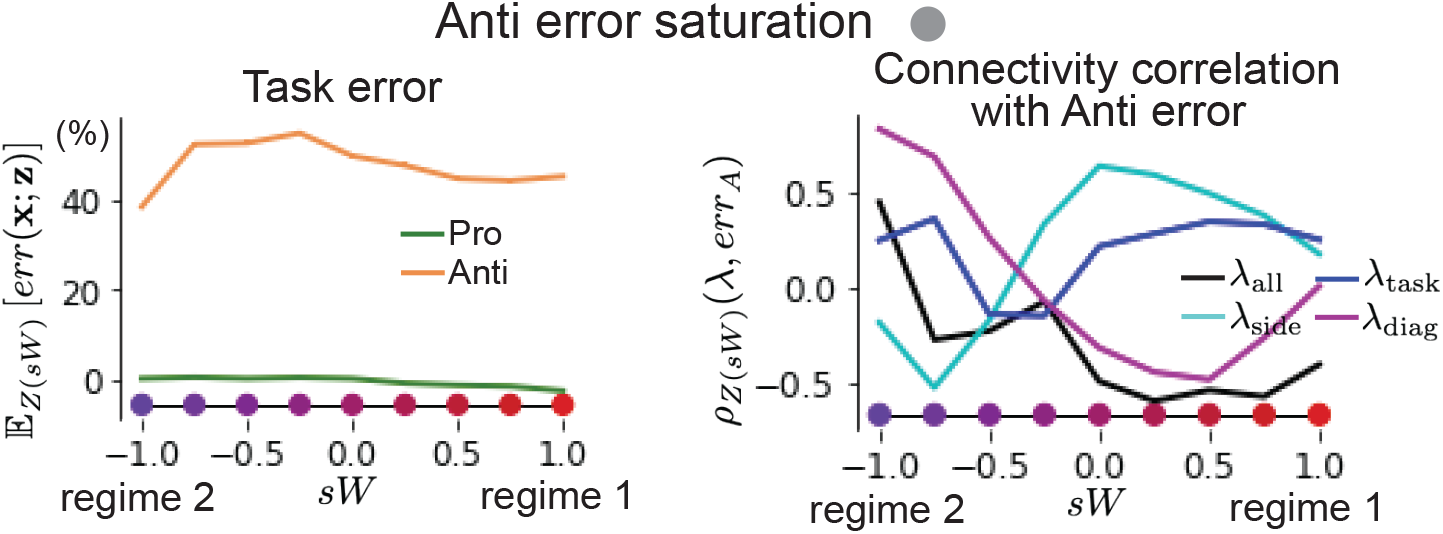
(Left) Mean and standard error of Pro and Anti error from regime 1 to regime 2 at *γ* = 0.85. (Right) Correlations of connectivity eigenvalues with Anti error from regime 1 to regime 2 at *γ* = 0.85.

#### 5.5.3 EPI details for the SC model

To write the emergent properties of Equation 9 in terms of the EPI optimization, we have

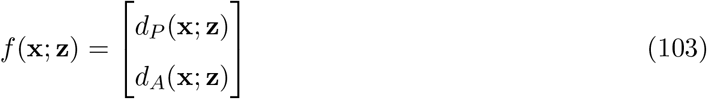

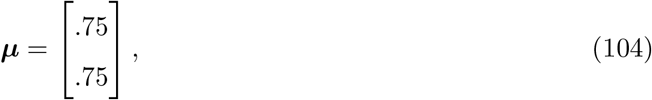

and

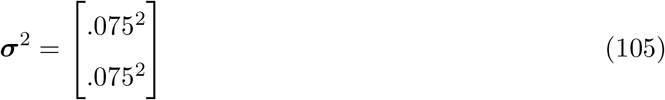

(see Sections 5.1.3–5.1.4, and example in Section 5.1.5).

**Figure S16:**
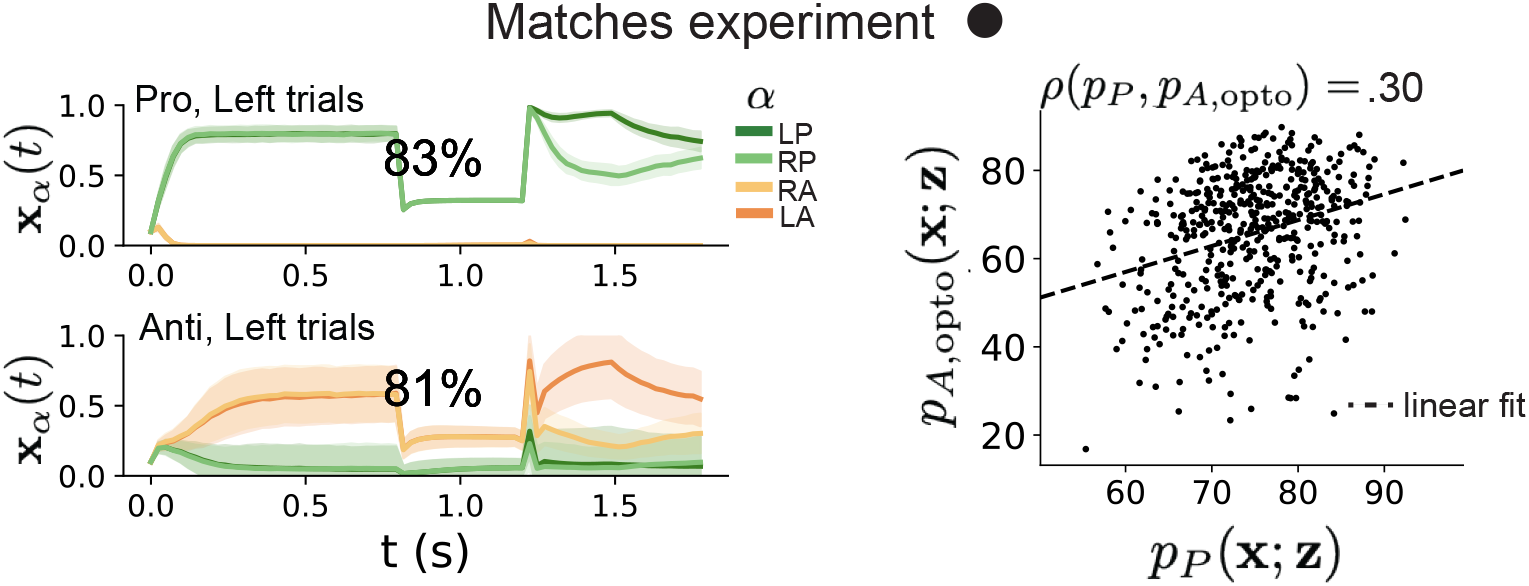
(Left) Mean and standard deviation (shading) of responses of the SC model at the mode of the EPI distribution to delay period inactivation at *γ* = 0.675. Accuracy in Pro (top) and Anti (bottom) task is shown as a percentage. (Right) Anti accuracy following delay period inactivation at *γ* = 0.675 versus accuracy in the Pro task across connectivities in the EPI distribution.

**Figure S17:**
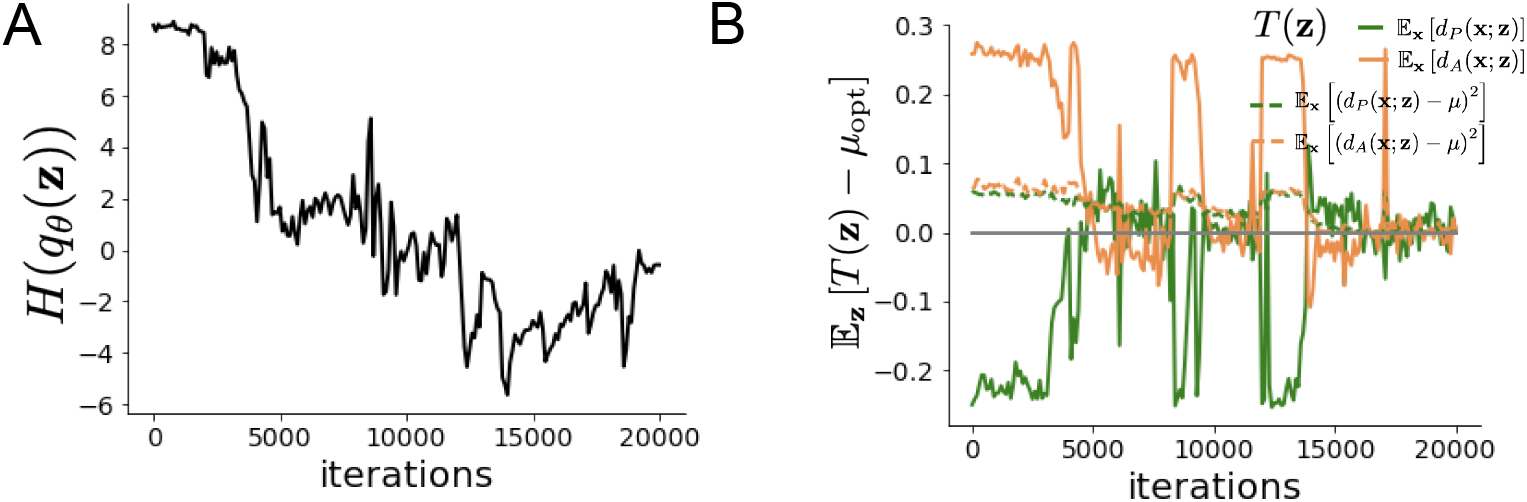
EPI optimization of the SC model producing rapid task switching. **A**. Entropy throughout optimization. **B**. The emergent property statistic means and variances converge to their constraints at 20,000 iterations following the tenth augmented lagrangian epoch.

Throughout optimization, the augmented lagrangian parameters *η* and *c*, were updated after each epoch of *i*_max_ = 2, 000 iterations (see Section 5.1.4). The optimization converged after ten epochs (Fig. S16).

For EPI in Fig. 4C, we used a real NVP architecture with three coupling layers of affine transformations parameterized by two-layer neural networks of 50 units per layer. The initial distribution was a standard isotropic gaussian 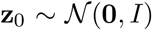 mapped to a support of **z**_*i*_ ∈ [−5, 5]. We used an augmented lagrangian coefficient of *c*_0_ = 10^2^, a batch size *n* = 100, and *β* = 2. The distribution was the greatest EPI distribution to converge across 5 random seeds with criteria *N*_test_ = 25.

**Figure S18:**
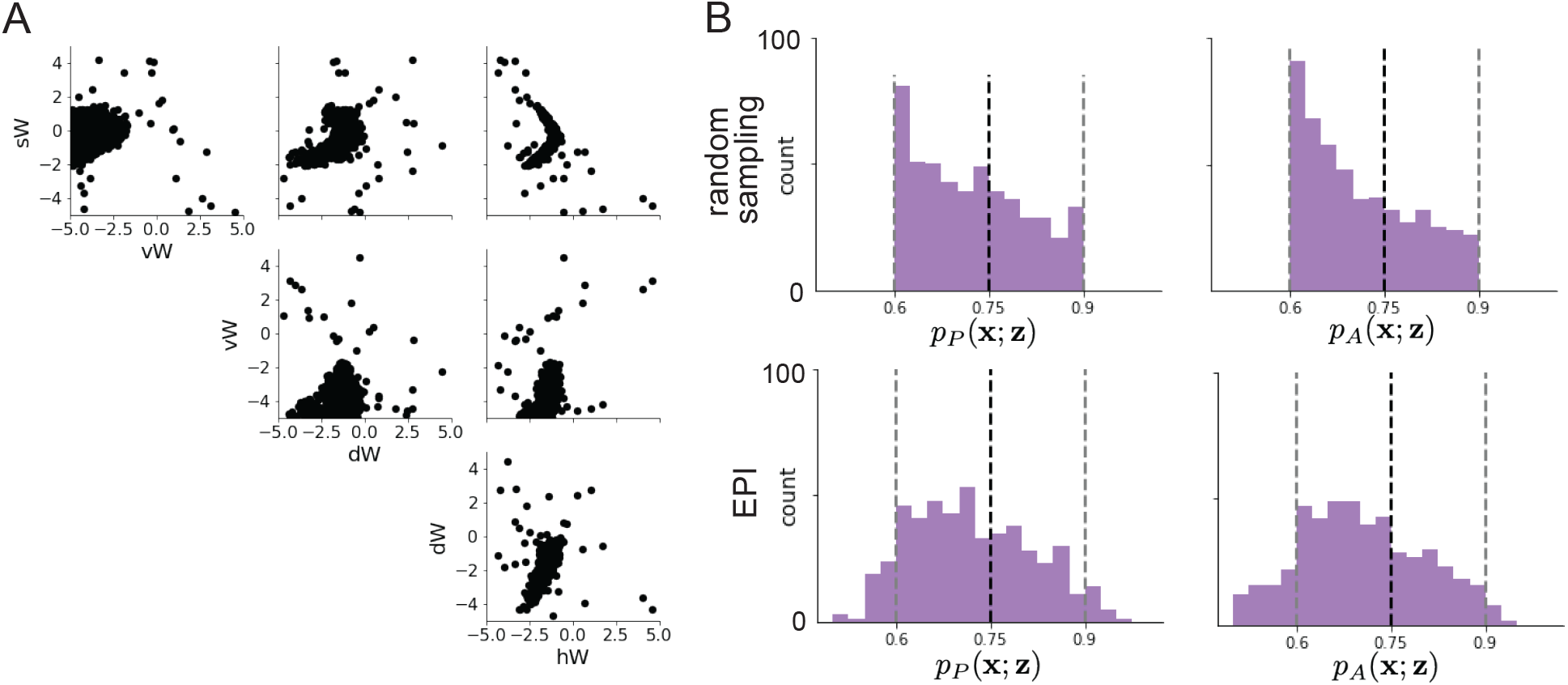
**A**.Rapid task switching SC connectivities obtained from random sampling. **B**. Task accuracies of the inferred distributions from random sampling (top) and EPI (bottom).

The bend in the EPI distribution is not a spurious result of the EPI optimization. The structure discovered by EPI matches the shape of the set of points returned from brute-force random sampling (Fig. S18A) These connectivities were sampled from a uniform distribution over the range of each connectivity parameter, and all parameters producing accuracy in each task within the range of 60% to 90% were kept. This set of connectivities will not match the distribution of EPI exactly, since it is not conditioned on the emergent property. For example the parameter set returned by the brute-force search is biased towards lower accuracies (Fig. S18B).

#### 5.5.4 Mode identification with EPI

We found one mode of the EPI distribution for fixed values of *sW* from 1 to −1 in steps of 0.25. To begin, we chose an initial parameter value from 500 parameter samples 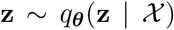 that had closest *sW* value to 1. We then optimized this estimate of the mode (for fixed *sW*) using probability gradients of the deep probability distribution for 500 steps of gradient ascent with a learning rate of 5 × 10^*−*3^. The next mode (at *sW* = 0.75) was found using the previous mode as the initialization. This and all subsequent optimizations used 200 steps of gradient ascent with a learning rate of 1 × 10^*−*3^, except at *sW* = −1 where a learning rate of 5 × 10^*−*4^ was used. During all mode identification optimizations, the learning rate was reduced by half (decay = 0.5) after every 100 iterations.

#### 5.5.5 Sample grouping by mode

For the analyses in Figure 5C and Figure S15, we obtained parameters for each step along the continuum between regimes 1 and 2 by sampling from the EPI distribution. Each sample was assigned to the closest mode **z**\* (*sW*). Sampling continued until 500 samples were assigned to each mode, which took 2.67 seconds (5.34ms/sample-per-mode). It took 9.59 minutes to obtain just 5 samples for each mode with brute force sampling requiring accuracies between 60% and 90% in each task (115s/sample-per-mode). This corresponds to a sampling speed increase of roughly 21,500 once the EPI distribution has been learned.

#### 5.5.6 Sensitivity analysis

At each mode, we measure the sensitivity dimension (that of most negative eigenvalue in the Hessian of the EPI distribution) **v**_1_(**z**\*). To resolve sign degeneracy in eigenvectors, we chose **v**_1_(**z**\*) to have negative element in *hW*. This tells us what parameter combination rapid task switching is most sensitive to at this parameter choice in the regime.

#### 5.5.7 Connectivity eigendecomposition and processing modes

To understand the connectivity mechanisms governing task accuracy, we took the eigendecomposition of the connectivity matrices *W* = *Q*Λ*Q*^−1^, which results in the same eigenmodes **q**_*i*_ for all *W* parameterized by **z** (Fig. S14A). These eigenvectors are always the same, because the connectivity matrix is symmetric and the model also assumes symmetry across hemispheres, but the eigenvalues of connectivity (or degree of eigenmode amplification) change with **z**. These basis vectors have intuitive roles in processing for this task, and are accordingly named the *all* eigenmode - all neurons co-fluctuate, *side* eigenmode - one side dominates the other, *task* eigenmode - the Pro or Anti populations dominate the other, and *diag* mode - Pro- and Anti-populations of opposite hemispheres dominate the opposite pair. Due to the parametric structure of the connectivity matrix, the parameters **z** are a linear function of the eigenvalues ***λ*** = [*λ*_all_, *λ*_side_, *λ*_task_*λ*_diag_]^⊤^ associated with these eigenmodes.

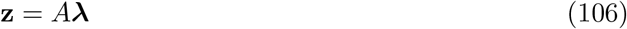

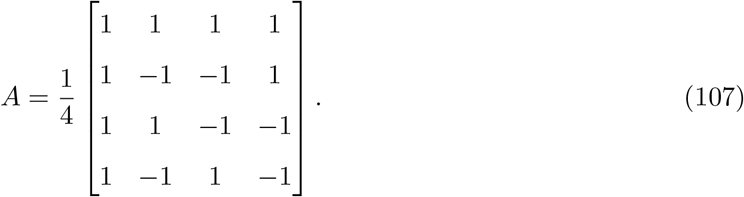

We are interested in the effect of raising or lowering the amplification of each eigenmode in the connectivity matrix by perturbing individual eigenvalues *λ*. To test this, we calculate the unit vector of changes in the connectivity **z** that result from a change in the associated eigenvalues

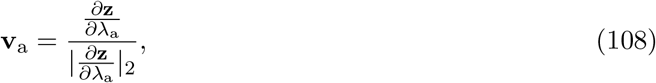

where

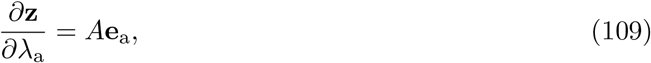

and e.g. **e**_all_ = [1, 0, 0, 0]^⊤^. So **v**_a_ is the normalized column of A corresponding to eigenmode *a*. The parameter dimension **v**_*a*_ (*a* ∈ {all, side, task, and diag}) that increases the eigenvalue of connectivity *λ_a_* is **z**-invariant (Equation 109) and **v**_*a*_ ⊥ **v**_*b*≠*a*_. By perturbing **z** along **v**_*a*_, we can examine how model function changes by directly modulating the connectivity amplification of specific eigenmodes, which having interpretable roles in processing in each task.

#### 5.5.8 Modeling optogenetic silencing

We tested whether the inferred SC model connectivities could reproduce experimental effects of optogenetic inactivation in rats [48]. During periods of simulated optogenetic inactivation, activity was decreased proportional to the optogenetic strength *γ* ∈ [0, 1]

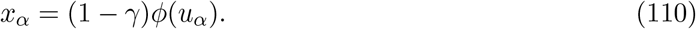

Delay period inactivation was from 0.8 < *t* < 1.2.

